# An autonomous oscillator times and executes centriole biogenesis

**DOI:** 10.1101/510875

**Authors:** Mustafa G. Aydogan, Thomas L. Steinacker, Mohammad Mofatteh, Lisa Gartenmann, Alan Wainman, Saroj Saurya, Siu S. Wong, Felix Y. Zhou, Michael A. Boemo, Jordan W. Raff

## Abstract

The accurate timing and execution of organelle biogenesis is crucial for cell physiology. Centriole biogenesis is regulated by Polo-like kinase 4 (Plk4) and initiates in S-phase when a daughter centriole grows from the side of a preexisting mother. Here we show that Plk4 forms an adaptive oscillator at the base of the growing centriole to initiate and time centriole biogenesis, ensuring that centrioles grow at the right time and to the right size. The Plk4 oscillator is normally entrained to the cell-cycle oscillator, but can run autonomously of it – explaining how centrioles can duplicate independently of cell cycle progression under certain conditions. Mathematical modelling indicates that this autonomously oscillating system is generated by a time-delayed negative-feedback loop in which Plk4 inactivates its centriolar receptor through multiple rounds of phosphorylation. We postulate that such organelle-specific autonomous oscillators could regulate the timing and execution of organelle biogenesis more generally.

## Introduction

Albert Claude’s landmark paper (Claude, 1943) challenged the idea that cells are a mere bag of enzymes whose contents grow freely in the cytoplasm with no active regulation. We now appreciate the diverse and compact nature of the many organelles in the cytoplasm (Marsh et al., 2001), yet the physical mechanisms that regulate the number and size of these organelles remain largely unknown (Marshall, 2016). For most organelles in the cell, however, this question has been difficult to address, as the variation in their numbers and 3D-shape has made it challenging to monitor their growth—or to even determine which parameter (e.g. their surface area, volume, or perhaps the amount of a limiting component) best defines their size.

Centrioles are highly structured organelles that form centrosomes and cilia (Bettencourt-Dias et al., 2011; Nigg and Holland, 2018; Nigg and Raff, 2009). Their linear structure and tightly controlled pattern of duplication makes them an attractive model with which to study organelle biogenesis (Goehring and Hyman, 2012; Marshall, 2016). Most cells are born with a single pair of centrioles that duplicate precisely once during S-phase, when a daughter centriole grows out orthogonally from the base of each mother until it reaches the same size as its mother (Banterle and Gönczy, 2017; Fırat-Karalar and Stearns, 2014; Nigg and Holland, 2018). To monitor the dynamics of centriole growth, we recently examined living syncytial *Drosophila* embryos where we could follow the assembly of hundreds of centrioles as they duplicate in near-synchrony in a common cytoplasm (Aydogan et al., 2018). These studies revealed that centriole growth in these embryos is homeostatic: when centrioles grow slowly, they grow for a longer period; when centrioles grow quickly, they grow for a shorter period. As a result, centrioles grow to a consistent size.

Polo-like kinase 4 (Plk4) is the master regulator of centriole biogenesis, and it is initially recruited to a ring around the mother centriole, but this ring ultimately resolves into a single focus on the side of the mother, defining the site of daughter centriole assembly (Banterle and Gönczy, 2017; Fırat-Karalar and Stearns, 2014; Leda et al., 2018; Nigg and Holland, 2018; Ohta et al., 2018). Unexpectedly, we found that Plk4 not only determines the position of this site, but also helps to establish the inverse relationship between the rate and period of daughter centriole growth (Aydogan et al., 2018). Plk4 presumably influences the rate of centriole growth by phosphorylating Ana2/STIL to promote its interaction with Sas-6 and so the assembly of the central cartwheel (Dzhindzhev et al., 2014; Kratz et al., 2015; Ohta et al., 2014), the 9-fold symmetric structure that forms the backbone of the growing daughter centriole (Kitagawa et al., 2011; van Breugel et al., 2011; 2014). It is less clear, however, how Plk4 might inversely adjust and determine the period of centriole growth.

Recent studies have shown that Plk4 localizes to centrioles in a cyclical manner in both fly embryos (Aydogan et al., 2018) and human cultured cells (Ohta et al., 2018), but the functional significance of this localisation pattern is unclear. Here, we demonstrate that Plk4 forms a cell-cycle autonomous oscillator at the base of the growing centriole that initiates and times centriole biogenesis in fly embryos.

## Results and Discussion

### Plk4 levels oscillate at the base of growing daughter centrioles

To investigate the cyclical recruitment of Plk4 to the centrioles, we generated flies transgenically expressing Plk4-mNeonGreen (Plk4-NG) under the control of its own promoter in a *Plk4* mutant background. We monitored centriolar Plk4-NG levels in living *Drosophila* syncytial embryos, where the duration of S-phase gradually elongates over nuclear cycles 11-13 (Figures 1A, S1 and S2A; Video S1). Centriolar Plk4-NG levels oscillated during each nuclear cycle: the oscillations started in mid-late mitosis, peaked in early-mid S-phase, and were minimal in early mitosis (Figures 1A and S2A). We used mathematical regression to fit these oscillations (Figures S1C and S1D), focusing our analysis on S-phase, the period when centrioles are growing (Figure 1B). The Plk4 oscillations exhibited adaptive behaviour: their amplitude (*A*) dampened during successive cycles, but their period (*T*)—as judged by the full width at half maximum amplitude—lengthened, so that the total amount of Plk4 recruited to centrioles (i.e., the area under the entire S-phase oscillation curve, *Ω*) remained relatively constant over each cycle (Figure 1C).

**Figure 1.**
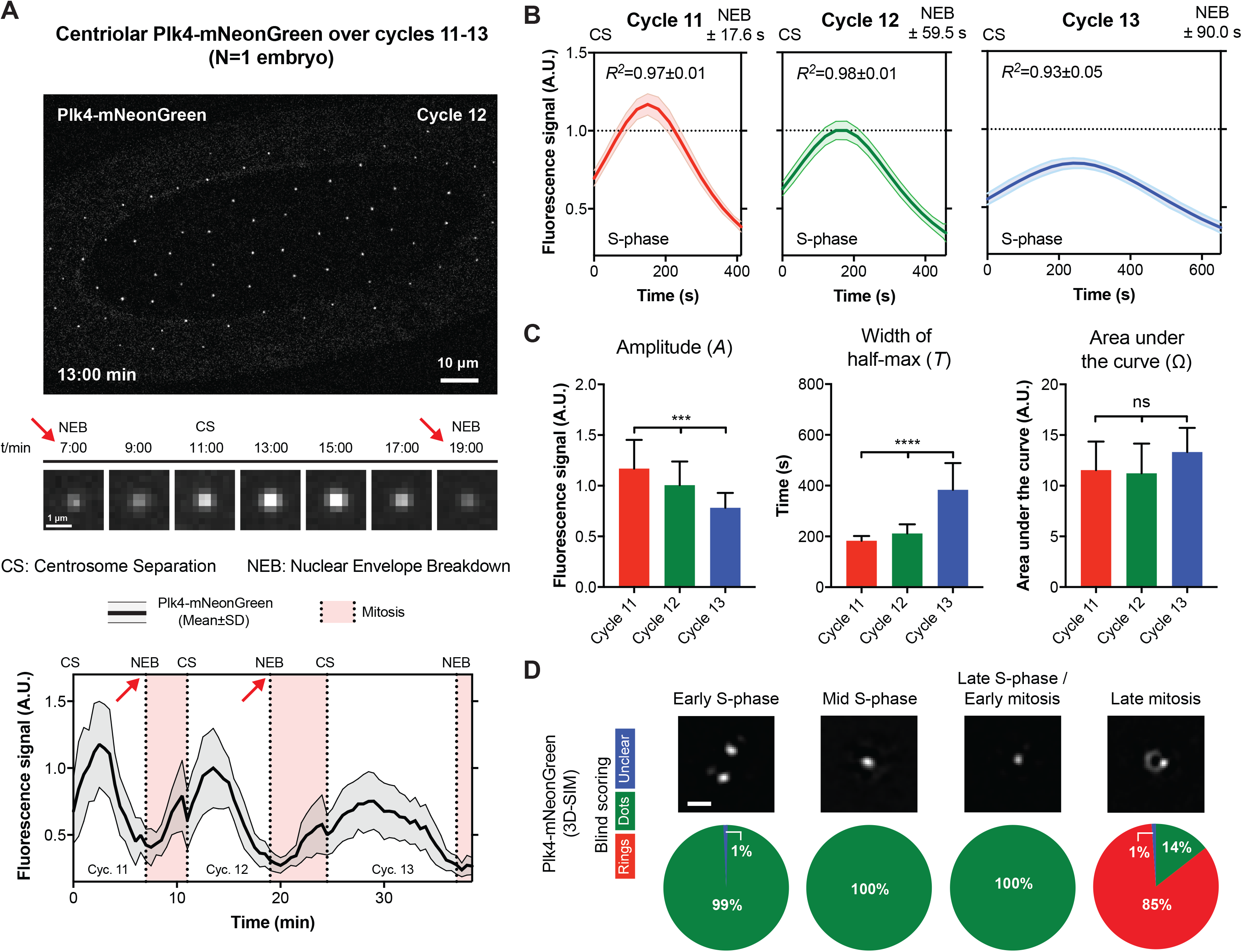
Plk4 levels oscillate at the centriole. **(A)** (Top panel) Micrograph shows an image from a time-lapse movie of an embryo expressing Plk4-NG. (Middle panels) Micrographs illustrate the centriolar Plk4-NG oscillation during nuclear cycle 12—obtained by superimposing all the Plk4-NG foci at each timepoint (n=60) (see *Materials and Methods*). (Bottom panel) Quantification of centriolar Plk4-NG levels during nuclear cycles 11-13 in a single embryo (*red* arrows highlight equivalent timepoints in the middle panels). **(B)** Graphs show the mathematical regression of centriolar Plk4-NG dynamics during S-phase of cycles 11-13 (Regression Mean±SEM). *R^2^* values indicate goodness-of-fit. N≥15 embryos; n=~24, 37, and 53 centrioles (mean) per embryo over cycles 11-13, respectively. **(C)** The bar charts quantify the oscillation parameters – derived from the data shown in (B). Data are presented as Mean±SD. Statistical significance was assessed using an ordinary one-way ANOVA test (for Gaussian-distributed data) or a Kruskal-Wallis test (***, P<0.001; ****, P<0.0001; ns, not significant). **(D)** Micrographs show, and piecharts quantify, the distribution of Plk4-NG at centrioles assessed by 3D-SIM at the indicated phases of the nuclear cycle (see *Materials and Methods*). N=6 embryos per cell cycle stage; n=20 centrioles per embryo; all images were scored blindly by 3 assessors and the mean score is shown (Scale bar = 0.5 μm). See also Figures S1 and S2A.

Plk4 is initially recruited to a ring around the mother centriole that resolves into a single hub that defines the site of daughter centriole assembly (Banterle and Gönczy, 2017; Firat-Karalar and Stearns, 2014; Nigg and Holland, 2018). To examine how this localisation is spatially linked to the Plk4 oscillations, we used 3D-Structured Illumination super-resolution Microscopy (3D-SIM) to assess the centriolar localisation of Plk4 during the nuclear cycles in living embryos. Plk4-NG was only briefly detectable in a ring during late-mitosis; at all other stages it appeared as a single hub (Figure 1D). We conclude that, during S-phase, centriolar Plk4-NG levels oscillate primarily at the base of the growing daughter centriole.

### Plk4 oscillations time and execute centriole biogenesis

To test whether the Plk4 oscillations were important for centriole biogenesis, we generated flies co-expressing Plk4-NG (in a *Plk4* mutant background) and the centriole cartwheel component Sas-6-mCherry, which is incorporated into the base of the growing daughter centriole cartwheel and can be used to monitor centriole growth in fly embryos (Aydogan et al., 2018). These flies laid embryos that often failed to hatch (Figure S3C), but we simultaneously measured Plk4 oscillations and centriole growth in the embryos that developed normally (Figures 2A, S3A and S3B; Video S2). The mother centrioles in these embryos were often slightly delayed in initiating daughter centriole growth (Figures 2A, S3D and S3E), allowing us to measure the amount of Plk4 at the centrioles when daughter centrioles either started or stopped growing (*coloured dotted* lines; Figure 2A).

**Figure 2.**
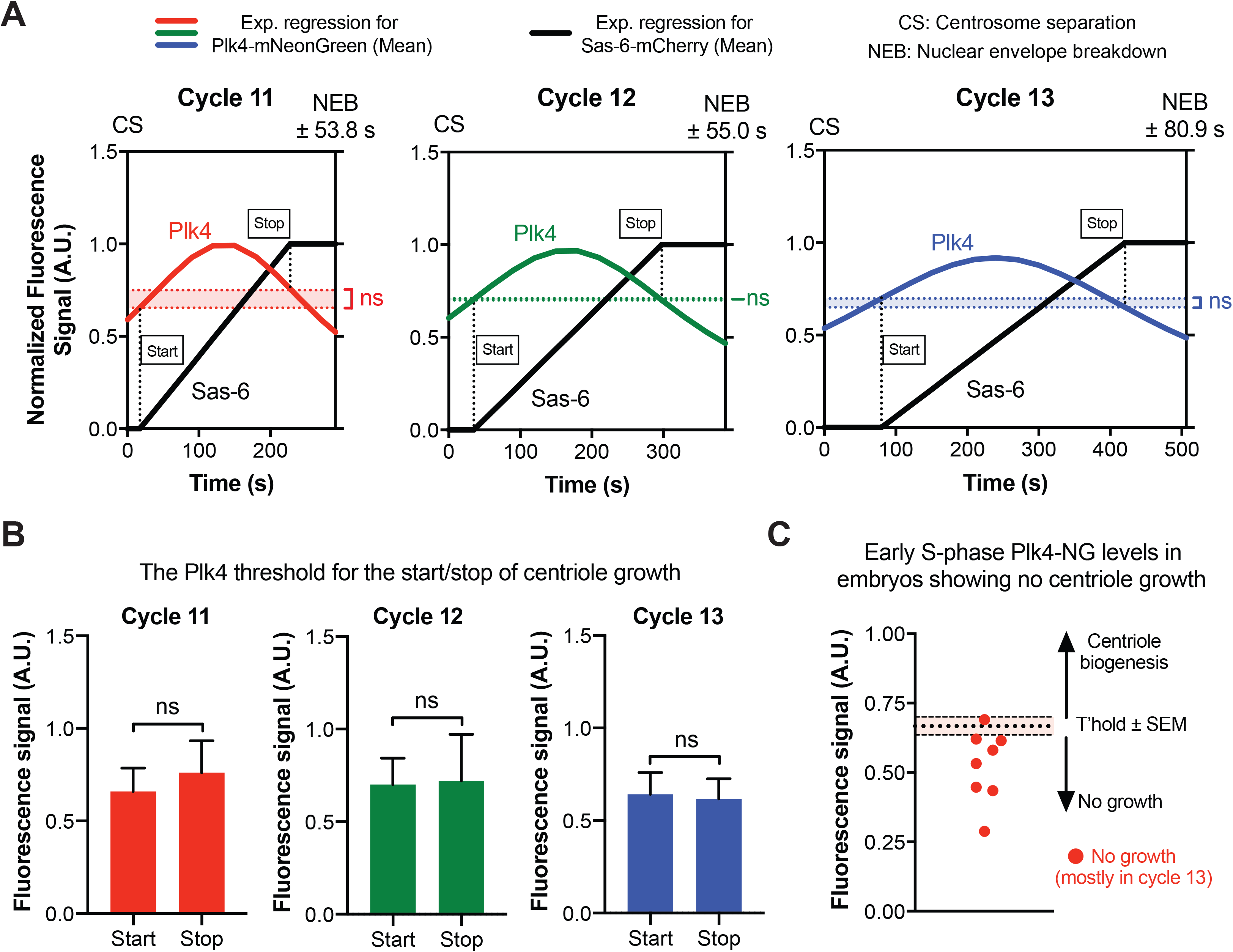
Plk4 oscillations initiate and time centriole biogenesis. **(A)** Graphs show the mean regression of Plk4-NG oscillations (*red, green* and *blue* lines for cycles 11-13, respectively) and centriole growth (monitored by Sas-6-mCherry incorporation—*black* lines) measured simultaneously in embryos during S-phase of cycles 11-13. For ease of presentation, the SEM for these data are not shown, but are presented in Figures S3A and S3B. Dotted lines indicate the centriolar Plk4 levels at which centrioles “start” or “stop” growing. N=17 embryos (cycles 11 and 12), and 8 embryos (cycle 13); n=19, 31, and 45 centrioles (mean) per embryo in cycle 11-13, respectively. See *Materials and Methods* for an explanation of data normalisation and scaling. **(B)** Bar charts quantify the centriolar Plk4-NG threshold levels at which centrioles start and stop growing during cycles 11-13 – derived from the data shown in (A). Data are presented as Mean±SD. Statistical significance was assessed using an unpaired *t* test with Welch’s correction (for Gaussian-distributed data) or an unpaired Mann-Whitney test (ns, not significant). **(C)** Eight embryos in which the centrioles did not grow (Figures S3D and S3E) were excluded from the analysis shown in (A) and (B); the scatter graph shown here illustrates how the mean amplitude of the Plk4 oscillations in each of these eight embryos (*red* dots) tended to be lower than the mean amplitude (±SEM) of the Plk4 oscillations in the embryos where the centrioles did grow. See also Figure S3.

Strikingly, the centriolar levels of Plk4 at which centriole growth initiated at each cycle (“*Start*”; Figures 2A and 2B) were not significantly different than the levels at which centriole growth stopped (“*Stop*”; Figures 2A and 2B). This suggests that there is a threshold level of centriolar Plk4 at each cycle that is required to support centriole growth: above this threshold the centrioles can grow, below this threshold they cannot. Although the threshold in these embryos was similar at successive nuclear cycles (Figure 2B), it seems that the absolute threshold level required to promote growth can vary depending on the conditions (see the *Plk4-NG* and *asl* “half-dose” experiments, below).

If the Plk4 threshold concept is correct, then mother centrioles that failed to recruit threshold levels of Plk4 would be predicted to not grow a daughter. We observed that the centrioles in a fraction of these embryos (mostly at nuclear cycle 13) separated at the start of S-phase but did not detectably incorporate Sas-6-mCherry, indicating that daughter centrioles did not grow (Figures S3D and S3E) – a defect that may explain why many of these embryos failed to develop properly (Figure S3C). Interestingly, centriolar Plk4 levels continued to oscillate in these embryos, but the average amplitude of these oscillations was lower than in the embryos in which centrioles continued to duplicate – and it was almost always below the average threshold at which centriole growth was normally initiated (Figure 2C). Taken together, these results strongly suggest that the Plk4 oscillations initiate, and then determine the duration of, centriole growth.

### Plk4 oscillations are normally entrained by the Cdk/Cyclin Oscillator

We next investigated whether the Plk4 oscillations were entrained by the core <u>C</u>dk/<u>C</u>yclin <u>o</u>scillator (CCO), which drives cell cycle progression in most eukaryotic cells. This appeared to be the case, as there was a strong positive correlation between the time at which the Plk4 oscillations peak and the length of S-phase (Figure 3A; Pearson *r* =0.8668, *P*<0.0001). Moreover, when we manipulated S-phase length – by halving the genetic dose of *Cyclin B* to elongate S-phase (Jacobs et al., 1998) (*CycB^1/2^* embryos), or halving the genetic dose of *Chk1* (*grapes*) to shorten S-phase (Sibon et al., 1997) (*grp^1/2^* embryos) – the phase of the Plk4 oscillations shifted accordingly (Figures 3B and 3C). We conclude that, under normal conditions, the Plk4 oscillator is entrained by the CCO.

**Figure 3.**
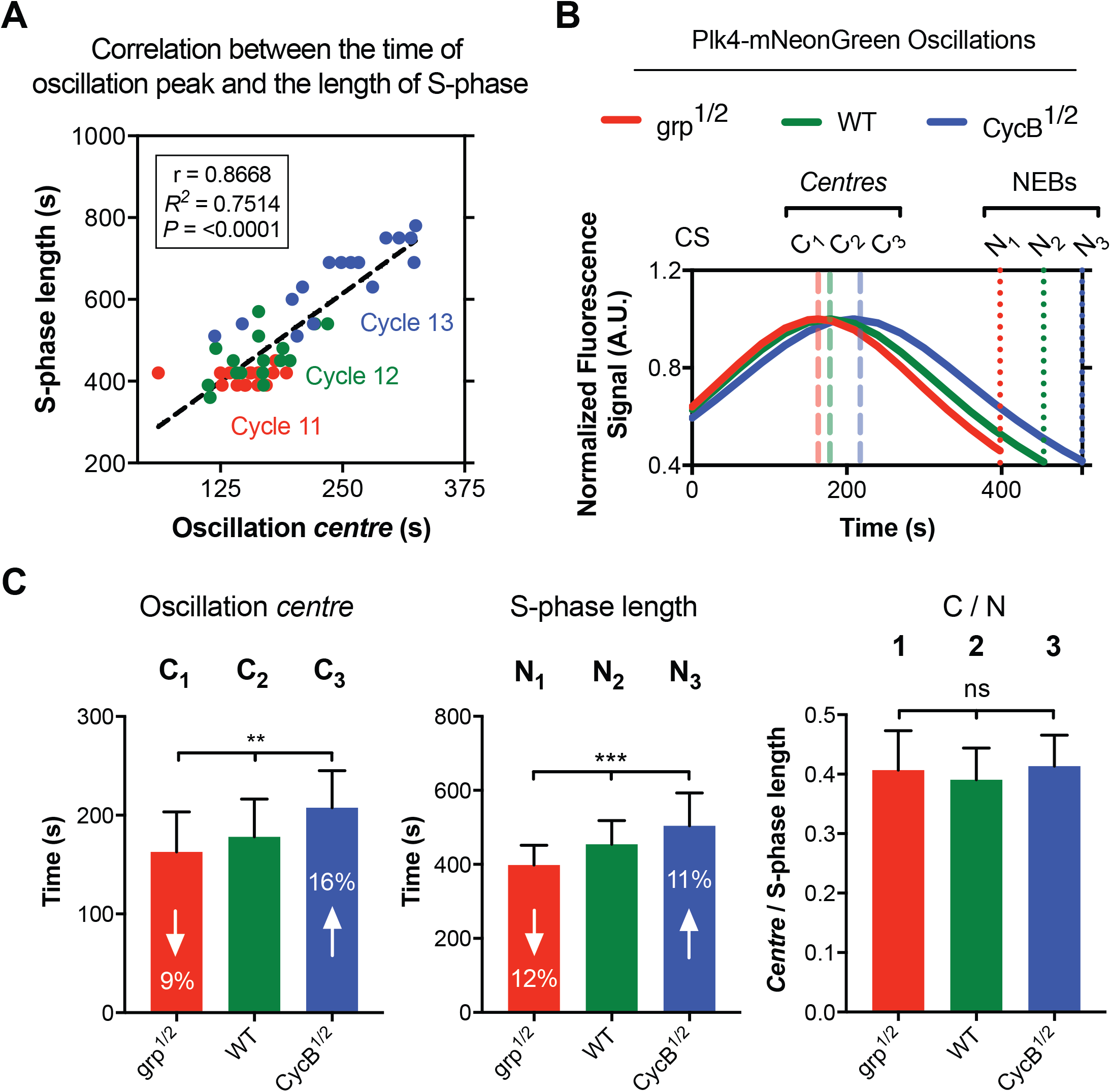
The Plk4 oscillator is entrained by the CCO. **(A)** Scatter plot shows the strong positive correlation between S-phase length and the centre (peak) of the Plk4-NG oscillations in cycles 11-13 (colour coded *red, green* and *blue*, respectively). The plot was generated with the data obtained from Figure 1B and was regressed using the line function in Prism 7 (GraphPad). Correlation strength was examined using Pearson’s correlation coefficient (r<0.40 weak; 0.40<r<0.60 moderate; r>0.6 strong); significance of correlation was determined by the p-value (P<0.05). **(B)** Graph shows the mean regression of Plk4-NG oscillations in nuclear cycle 12 of WT (*green*), *CycB^1/2^* (*blue*) and *grp^1/2^* (*red*) embryos. *Dashed* lines mark the *centre* (peak) of the Plk4-NG oscillations (denoted with C) and, *dotted* lines indicate the time of NEB (denoted with N) for each genotype. N≥14 embryos for each condition; n=55, 43, and 44 centrioles (mean) per embryo in WT, *CycB^1/2^* and *grp^1/2^* embryos respectively. To clearly illustrate the phase shift in the oscillations, the highest mean fluorescence signal for each group was normalised to 1. **(C)** Bar charts quantify the time at which the Plk4-NG oscillations peaked, the length of S-phase, and the ratio between them (C/N) – derived from the data shown in (B). Data are presented as Mean±SD. Statistical significance was assessed using an ordinary one-way ANOVA test (for Gaussian-distributed data) or a Kruskal-Wallis test (**, P<0.01; ***, P<0.001; ns, not significant).

### Plk4 forms an autonomous oscillator that can execute centriole duplication independently of a robust cell-cycle oscillator

It has long been known that centrioles can continue to duplicate in many systems even when several other aspects of cell cycle progression have been blocked (Balczon et al., 1995; Gard et al., 1990; McCleland and O’Farrell, 2008; Sluder et al., 1990). We wondered whether this might reflect an innate ability of Plk4 to continue to oscillate at centrioles and drive centriole biogenesis even if the CCO is perturbed. To test this possibility, we injected embryos with double-stranded RNAs (dsRNAs) targeting the three embryonic mitotic cyclins: A, B and B3. This treatment arrests embryos in an interphase-like state with intact nuclei that neither duplicate their DNA nor divide, but where centrosomes can continue to duplicate (McCleland and O’Farrell, 2008). We initially injected embryos in nuclear cycle 7-8 and monitored Plk4-NG behaviour ~30 min later, aiming to examine centriole duplication in embryos that went through a final round of mitosis before arresting in an interphase-like state (as judged by the lack of NEB). In all such embryos examined, we observed additional round(s) of centriole duplication without NEB (Figures 4A and S2B; Video S3). These duplication events were less synchronous than normal, and we observed occasional cases of overduplication – as seen previously when Cdk1 activity is inhibited in flies (Vidwans et al., 2003) – but a clear Plk4-NG oscillation was associated with each additional round of centriole duplication (*dotted* lines; Figures 4A and S2B).

**Figure 4.**
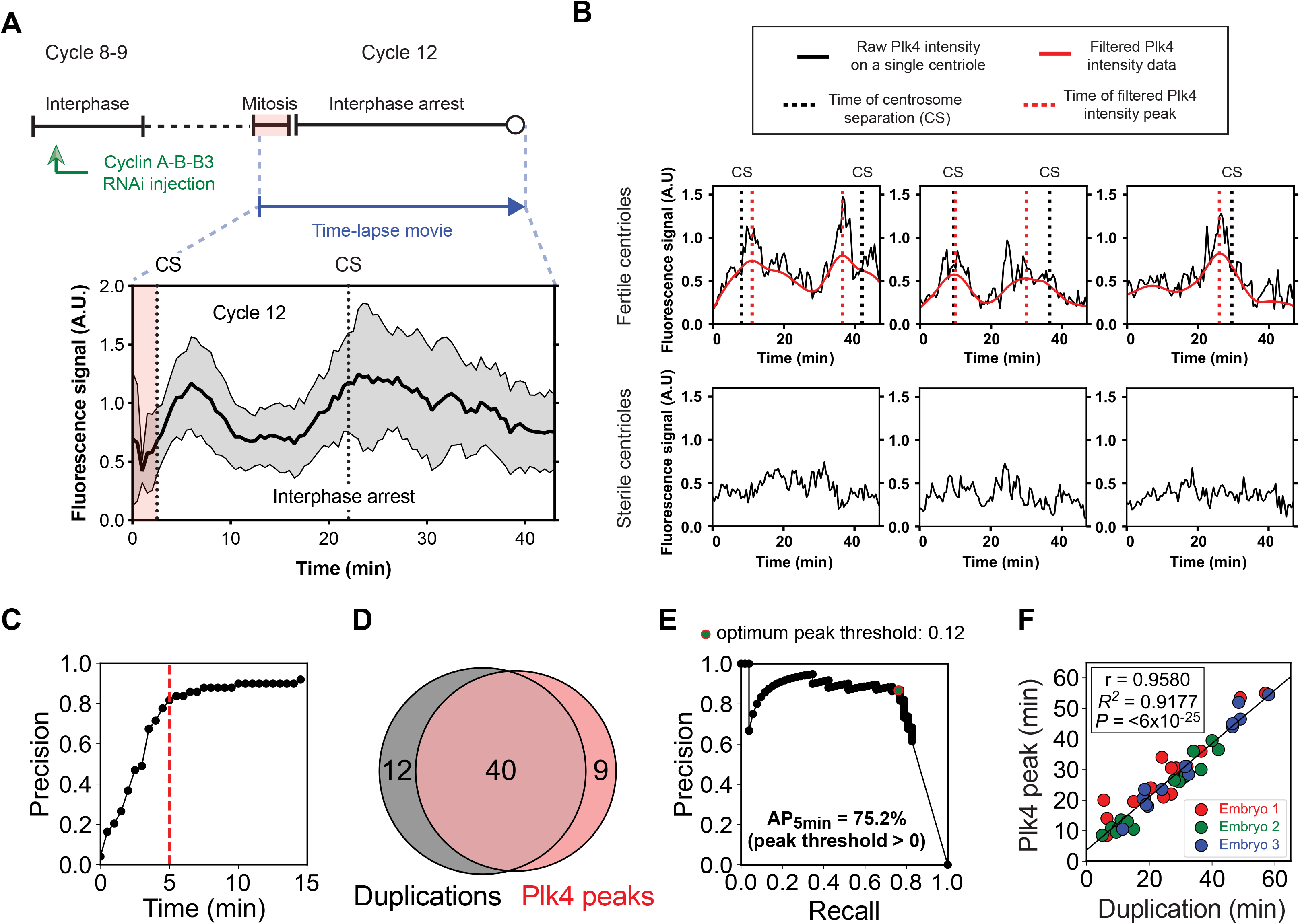
Plk4 levels can continue to oscillate and promote centriole duplication even when the CCO is perturbed. **(A)** Graph shows the Plk4-NG oscillations in an embryo injected with dsRNA against cyclin A-B-B3; the schema above the graph illustrates the experimental protocol. The nuclei in this embryo arrest in interphase, but centrioles go through an additional round of division—Centriole Separation (CS)—accompanied by a Plk4 oscillation. See Figure S2B for 5 additional examples; n=30 centrioles (mean) per embryo. **(B)** Graphs show the raw (*black* lines) and filtered (*red* lines) fluorescence intensity data of 3 individual “fertile” centrioles and 3 individual “sterile” centrioles within the same embryo. The fertile centrioles duplicate (*black dotted* lines), and these events were often closely associated with computed Plk4 oscillation peaks (*red dotted* lines). **(C)** An unbiased computational analysis of all 45 fertile centrioles in 3 embryos reveals that >80% of the computationally detected Plk4 oscillation peaks occur within 5 minutes of an experimentally observed duplication event. For comparison, simulation with randomly distributed centriole duplication events showed a mean time separation of 10.5 minutes (data not shown). **(D)** Venn diagram shows how, using this 5 minute window, the oscillation peaks can be used to predict duplication events with both high precision and high recall (40/49 Plk4 oscillation peaks are associated with a duplication event, and 40/52 duplication events are associated with a Plk4 oscillation peak). **(E)** Graph shows the ability of Plk4 oscillation peaks to ‘retrieve’ centriole duplication events across all peak prominences. All detected oscillation peaks were ranked in order of their peak prominence from high to low (*black* dots) and assigned uniquely to a duplication event if within a 5 min time window. The graph then plots the precision and recall values if the threshold for calling a peak were set as the peak prominence value of each peak (in descending order). Below the detected peak that is associated with a peak prominence threshold of 0.12, the precision dramatically drops, suggesting the existence of a minimum peak amplitude for centriole duplication. At this threshold, precision and recall are jointly optimised. Note, if there were no overall correlation between Plk4 peaks and a duplication event, the integrated area under the curve across all peak prominences or average precision (AP) for the 5 min time window (AP_5min_) would be ~50% (given by # duplications / (# duplications + # peaks)); so the score of ~75% indicates a meaningful correlation. **(F)** Graph shows the significant correlation between the timing of the computationally determined Plk4 peaks and the respective, temporally closest duplication events that were experimentally observed. Correlation strength was examined using Pearson’s correlation coefficient (r<0.40 weak; 0.40<r<0.60 moderate; r>0.6 strong); significance of correlation was determined by the p-value (P<0.05) (see *Materials and Methods* for a full description of this analysis). See also Figures S4 and S5.

We next injected embryos at an earlier stage (nuclear cycles 2-4) and monitored them ~90 min later. These embryos contained many centrioles that were dissociated from the non-dividing nuclei (Video S4). Centrioles in these embryos continued to duplicate, but in a stochastic manner: some centrioles were “fertile”, and duplicated one or more times, while others were “sterile” and did not duplicate at all (Figure S4A; Video S4). We therefore measured Plk4-NG fluorescence levels at individual centrioles. As expected, the raw intensity data was noisy (*black* lines; Figure 4B), but fertile centrioles appeared to exhibit Plk4-NG oscillations, whereas sterile centrioles did not. In support of this interpretation, the average centriolar Plk4-NG fluorescence level (expressed as signal-to-noise ratio – SNR) was significantly higher at fertile centrioles (Figure S4B), and the SNR values could distinguish (Figure S4C) and correctly predict whether a centriole is fertile or sterile ~74% and ~71% of the time, respectively (Figure S4D).

Upon filtering the raw oscillation data (*red* lines; Figure 4B), we found that the peaks of the Plk4-NG oscillations (*dotted red* lines; Figure 4B) were often associated with centriole duplication events (*dotted black* lines; Figure 4B). To test whether this association was functionally significant, we performed an unbiased computational analysis of all the 45 fertile centrioles that we observed in 3 different embryos (Figures 4C–E). This revealed that the centriolar Plk4-NG oscillation peaks predicted centriole duplication events with high precision (40/49 Plk4-NG peaks were associated with a duplication event that occurred within ±5 mins of the peak) and recall (40/52 duplication events occurred within ±5 mins of a Plk4-NG oscillation peak) (Figures 4C and 4D). Computer simulations revealed that a random distribution of the oscillation peaks and centriole duplication events lead to an average time of >10mins between the peaks and duplication events, indicating that the observed association was not random. Moreover, a rank ordering of the Plk4-NG oscillations based on their amplitude revealed that the higher the amplitude of the oscillation, the more likely it was to be associated with a centriole duplication event (Figure 4E), while plotting the relative timing of the Plk4-NG oscillations and the centriole duplication events revealed a strong positive correlation (Figure 4F; Pearson *r* =0.9580, *P*<0.0001). Together, these data strongly suggest that individual centrioles organise an autonomous Plk4 oscillator that can continue to drive stochastic centriole duplication events even in the absence of a robust CCO.

It is well established that centrioles can duplicate during S-phase but not during mitosis (Khodjakov et al., 2002; Zitouni et al., 2016). To test whether Plk4 oscillations could continue when embryos were arrested in mitosis, we injected embryos with the microtubule-depolymerising drug colchicine (Yuan and O’Farrell, 2015) (Figures S5A and S2C; Video S5). Some residual Plk4-NG oscillations were detectable in these embryos (Figures S5A and S2C), but their amplitude was dramatically reduced (Figures S5B and S2C). These Plk4 oscillations appeared to be sub-threshold for centriole biogenesis (Figure S5C), and we observed no further centriole duplication events in these embryos. Thus, the Plk4 oscillator appears to be calibrated by the CCO to ensure that it cannot execute centriole biogenesis during mitosis.

It is widely believed that the CCO acts primarily as a “ratchet” whose activity increases over the cell cycle to irreversibly trigger the sequential execution of cell cycle events such as DNA replication, centriole/spindle pole body duplication, and spindle assembly (Stern and Nurse, 1996; Swaffer et al., 2016; 2018). An interesting alternative possibility is that the CCO could act primarily as a “phase-locker” whose function is simply to entrain the phase of a network of autonomous oscillators, each of which is responsible for the timely execution of a specific cell-cycle event (Lu and Cross, 2010; Orlando et al., 2008; Rahi et al., 2016). Our observations are consistent with this latter model, and we propose that Plk4 forms such an autonomous oscillator, which is normally entrained by the CCO to execute centriole biogenesis.

### A mathematical model for the Plk4 oscillator in S-phase

How are Plk4 oscillations generated? Our observation that the Plk4 oscillator is adaptive – i.e., the *A* and *T* of the oscillations are inversely correlated to ensure a relatively constant *Ω* (Figures 1B and 1C) – suggests that centrioles can measure, or integrate, the total amount of Plk4 bound to them in each cell cycle. *Drosophila* Asterless (Asl) recruits Plk4 to centrioles and appears to activate it, which in turn allows Plk4 to phosphorylate itself and Asl at multiple sites (Boese et al., 2018; Dzhindzhev et al., 2010; Klebba et al., 2015). Moreover, human Asl (Cep152) also binds, and is phosphorylated by, Plk4 *in vitro* (Cizmecioglu et al., 2010; Hatch et al., 2010). Together, these observations suggested the following schema (Figures 5A and 5B): At the start of each oscillation cycle, Asl receptors, located at the base of mother centrioles, bind Plk4 with high affinity to recruit it to centrioles and thereby activate it (Figures 5A_(i)_ and B_(i)_). The activated Plk4 then phosphorylates itself (Cunha-Ferreira et al., 2013; Holland et al., 2010; Klebba et al., 2013) and Asl (Boese et al., 2018) at multiple sites, providing a monitoring mechanism for each Asl receptor to integrate how much Plk4 kinase activity it has experienced over time (Figures 5A_(ii)_ and B_(ii)_). The active, Asl-bound Plk4 also phosphorylates Ana2/STIL to promote centriole growth (Dzhindzhev et al., 2014; Kratz et al., 2015; Ohta et al., 2014) (Figure 5A_(ii)_) – explaining why a threshold level of centriolar Plk4 is required for centriole biogenesis – but this reaction is not important for the generation of Plk4 oscillations *per se*, so we do not consider it further. We propose that, once phosphorylated at multiple sites, the Asl receptor switches to a state that binds Plk4 with low-affinity, which is followed by the release, and potential degradation, of the phosphorylated Plk4 (Figures 5A_(iii)_ and B_(iii)_). This leaves behind the fully phosphorylated Asl-receptor that can no longer recruit Plk4, and so can no longer promote centriole growth. We envisage that this circuit is “re-set” during early mitosis by a phosphatase (whose centriolar activity presumably oscillates out of phase with the Plk4 oscillation) that dephosphorylates Asl (*red* arrow; Figure 5B) – although we do not consider this in our model.

**Figure 5.**
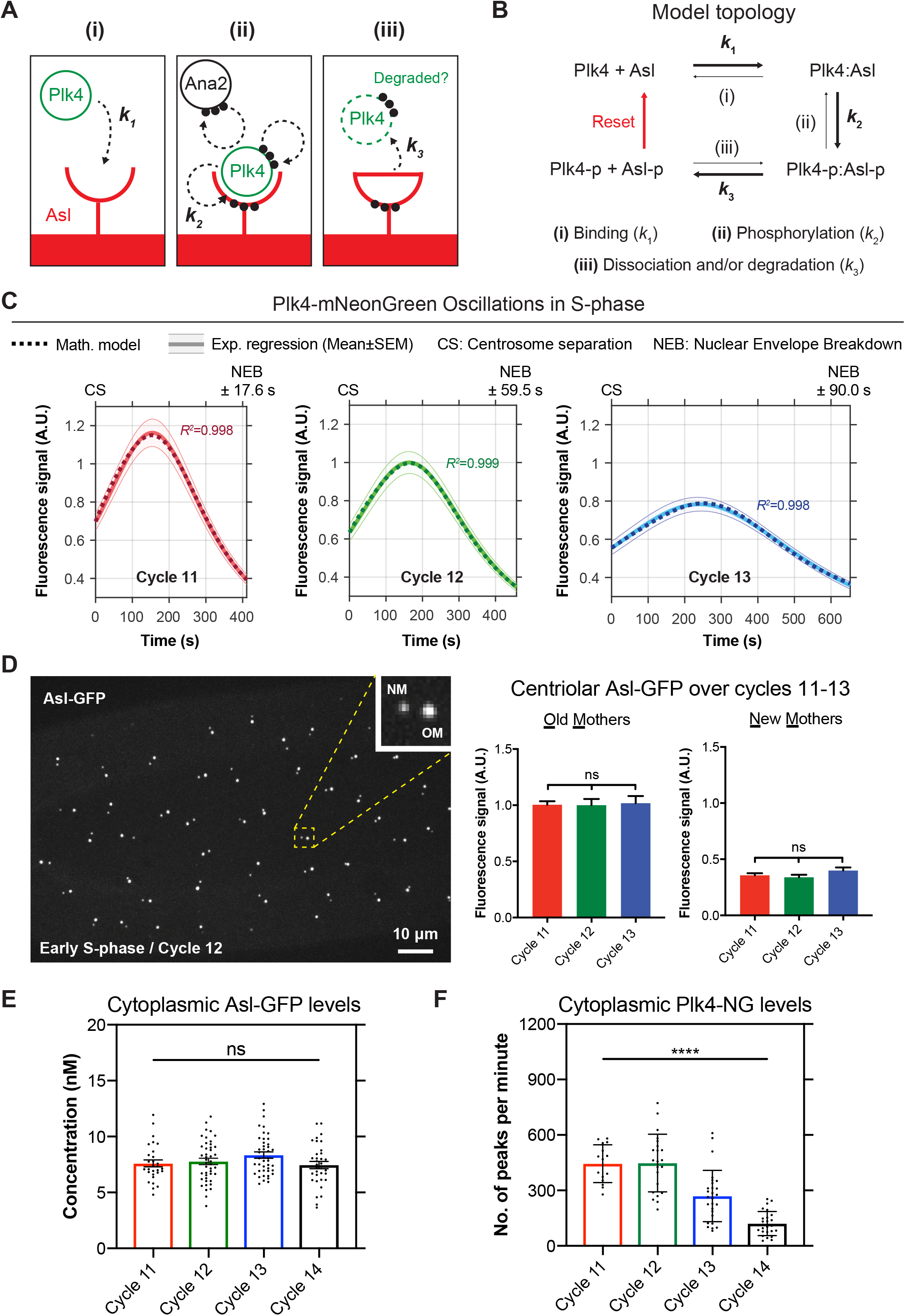
A mathematical model of the Plk4 oscillation, and experimental investigations to test its predictions. **(A)** Diagram of the model. **(i)** During mitosis, Asl receptors at the surface of the mother centriole start to bind Plk4 with high affinity (*k*_1_). **(ii)** Once bound, Plk4 is activated, and it starts to phosphorylate itself, Ana2 and Asl (*k*_2_) at multiple sites (*dotted black* arrows and *black* dots) (Boese et al., 2018; Dzhindzhev et al., 2010; Klebba et al., 2015). **(iii)** After several rounds of phosphorylation we speculate that Asl is converted to a state with low affinity for Plk4, so phosphorylated Plk4 is released (*k*_3_)—and likely degraded (Cunha-Ferreira et al., 2013; Holland et al., 2010; Klebba et al., 2013). These Asl receptors are now inactivated and can no longer promote centriole growth. **(B)** Schematic depicts the topology of the mathematical model (see *Materials and Methods* for full details of the model). Asl-p and Plk4-p indicate phosphorylated proteins. Bold arrows indicate the dominant direction of the reactions. We speculate that a phosphatase removes the phosphate groups from Asl during mitosis to reset the system for the next oscillation (*red* arrow). This step is not incorporated into our mathematical model, which discretely models centriolar Plk4-NG levels only during S-phase of each cycle. **(C)** Graphs show the Lorentzian fit of the Plk4-NG oscillation data during S-phase of cycles 11-13 (*solid* lines) overlaid with mathematical solutions to the model (*dotted* lines). *R^2^* values indicate goodness-of-fit for the mathematical solutions. **(D)** Micrograph shows Asl-GFP at newly separated old mother (OM) and new mother (NM) centrioles. Graphs quantify fluorescence levels on each centriole at the start of S-phase of each cycle. Asl preferentially localises to OM centrioles at the start of S-phase (Novak et al., 2014), but the amount of Asl at either OM or NM centrioles at the start of S-phase remains constant at successive nuclear cycles. N≥14 embryos for each cell cycle; n=48, 70, and 130 centrioles (mean) per embryo in cycle 11-13, respectively. Data are presented as Mean±SEM. Statistical significance was assessed using an ordinary one-way ANOVA test (for Gaussian-distributed data) or a Kruskal-Wallis test (ns, not significant). **(E and F)** Graphs show the cytosolic concentration of Asl-GFP at the start of each nuclear cycle measured by FCS (E) and the relative abundance of Plk4-NG at the start of each nuclear cycle measured by PeCoS (F) (see *Materials and Methods*; Figures S7 and S8). Every data point represents the average of 4-6 recordings from a single embryo or an individual 180 sec recording in (E) and (F), respectively. Statistical significance was assessed using an ordinary one-way ANOVA test (for Gaussian-distributed data) or a Kruskal-Wallis test (****, P<0.0001; ns, not significant). See also Figures S6–8 and Tables S1 and S7.

This model exemplifies a type of “time delayed negative-feedback” network where an inhibitor (Plk4) inhibits its own activator (Asl) (Novák and Tyson, 2008). This network maps onto a set of coupled linear ordinary differential equations (see the mathematical modelling section in *Materials and Methods*), which we solved analytically. This model fit the discrete Plk4 oscillation data from each S-phase of nuclear cycles 11-13 very well (Figure 5C; *R*^2^>0.99), generating solutions that were within a reasonable and generally narrow parameter space (Figures 5C and S6; Tables S1 and S7). Importantly, this model considers only the behaviour of Asl and Plk4; it is likely that other factors, such as Ana2/STIL, will modulate Asl and/or Plk4 behaviour at the centriole assembly site. This may help to explain, for example, how the initial ring of Plk4 at the centriole resolves into a single hub: we speculate that Plk4 may rapidly phosphorylate Asl receptors that are not at the site of centriole assembly, thus quickly inactivating them so that Plk4 only briefly localises to the ring. Additional factors present at the assembly site may stabilise the Asl:Plk4 interaction (perhaps by influencing the structural conformation of Asl in ways that dampen its ability to be phosphorylated, or by simply affecting Plk4’s kinase activity), so allowing Plk4 to persist at this site to promote cartwheel growth.

### Testing the mathematical model: centriolar Asl levels remain constant at successive nuclear cycles, whereas cytoplasmic Plk4 levels decrease

The parameters generated by our mathematical model made two testable predictions. First, centriolar Asl levels are predicted to remain relatively constant at each cycle (*A_tot_*; Table S1). We confirmed that this was the case at both old mother (OM) and new mother (NM) centrioles (Figure 5D) (see below), and *Fluorescence Correlation Spectroscopy* (FCS) revealed that the cytosolic concentration of Asl-GFP – that is known to influence the amount of Asl at the centrioles (Novak et al., 2014) – remained constant at the start of each nuclear cycle (Figures 5E and S7). Second, the most significant difference between the oscillation parameters at nuclear cycles 11-13 is the rate at which Plk4 binds to Asl (*k*_1_; Figures 5A and 5B), which is predicted to decrease at successive nuclear cycles (Table S1). This parameter depends on the cytoplasmic concentration of Plk4 that, unfortunately, was too low to be measured by conventional FCS. We therefore developed a new method, *Peak Counting Spectroscopy* (PeCoS), which can accurately measure relative protein abundance at much lower concentrations (see *Materials and Methods*) (Figure S8). This revealed that, unlike cytoplasmic Asl levels (Figure 5E), cytoplasmic levels of Plk4 tended to decrease at successive nuclear cycles (Figure 5F), as predicted by our model.

Interestingly, as reported previously (Novak et al., 2014; 2016), OM centrioles initially associate with more Asl than NM centrioles (Figure 5D), so it is unclear how the OM and NM both grow daughters of similar size when they associate with such different amounts of Asl at the start of S-phase. We speculate that Asl may initially be recruited rapidly to the site of daughter centriole assembly on the NM when the OM and NM centrioles first separate during mitosis – so that OM and NM centrioles have similar levels of Asl at the centriole assembly site when daughter centriole assembly is initiated in S-phase. Asl may then be recruited more gradually to the rest of the NM centriole (Novak et al., 2014).

### The Plk4 oscillator can adapt to changes in input (cytoplasmic Plk4 levels) to maintain a constant output (centriole size)

Our observations suggest a rationale for why centriole biogenesis may be regulated by an oscillatory system: in the model we describe here, Asl functions as an *integrator* (Ferrell, 2016; Somvanshi et al., 2015) whose levels are kept constant so that the oscillator can monitor changes in the input (in this case, cytoplasmic Plk4 levels) and respond accordingly by adapting the oscillation to maintain a constant output (in this case, centriole size). If this is correct, the Plk4 oscillator should be able to adapt to changes in Plk4 levels to maintain a consistent centriole size, but not to changes in Asl levels. To test this, we initially monitored Plk4-NG oscillations in embryos laid by mothers where we genetically halved the dose of either Plk4-NG (hereafter *Plk4-NG^1/2^* embryos) or *asl* (hereafter *asl^1/2^* embryos). Centrioles appeared to duplicate normally in both sets of embryos, but the Plk4 oscillation parameters were altered: In *Plk4-NG^1/2^* embryos, *A* decreased but there was a compensatory increase in T, so *Ω* remained relatively constant (Figures 6A and S9A); in *asl^1/2^* embryos, *A* decreased, but there was no compensatory change in *T*, so *Ω* decreased (Figures 6B, S9B and S9C). Our mathematical model fitted both sets of data well (*R^2^*>0.99), generating a reasonable range of parameters (Tables S2 and S3), several of which we validated experimentally (see the mathematical modelling section in *Materials and Methods*).

**Figure 6.**
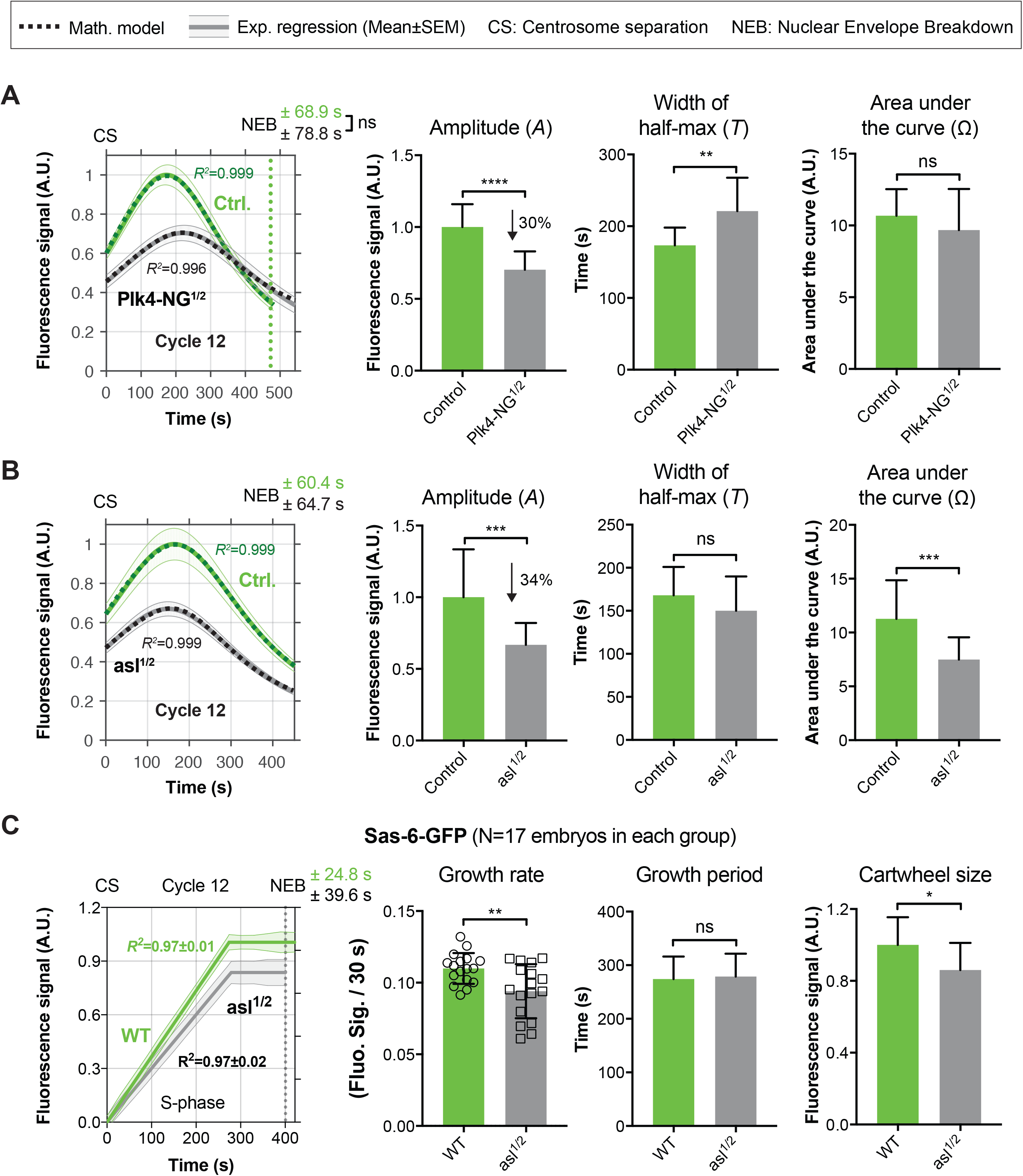
The Plk4 oscillator can adapt to changes in Plk4 concentration, but not to changes in Asl concentration. Graphs show the regression data (*solid* lines) and mathematical solutions (*dotted* lines) for Plk4-NG oscillations in cycle 12 for experiments where either the genetic dose of *Plk4-NG* was halved (*Plk4-NG^1/2^*), or **(B)** the genetic dose of *asl* was halved (*asl^1/2^*) (*grey* lines) compared to controls (*green* lines). In (A), N≥11 embryos for each condition; n=47 and 42 centrioles (mean) per embryo in *Control* or *Plk4-NG^1/2^* groups, respectively. In (B), N=18 embryos for each condition; n=44 and 43 centrioles (mean) per embryo in *Control* or *asl^1/2^* groups, respectively. Data are presented as Mean±SEM. Bar charts quantify oscillation parameters, as indicated; data are presented as Mean±SD. (C) Graph quantifies the parameters of cartwheel growth—as measured by Sas-6-GFP fluorescence incorporation (Aydogan et al., 2018)— in *WT* and *asl^1/2^* embryos; data are presented as Mean±SEM. Bar charts quantify growth parameters, as indicated; data are presented as Mean±SD. N=17 embryos for each condition; n=77 and 72 centrioles (mean) per embryo in WT or *asl^1/2^* groups, respectively. Statistical significance was assessed using an unpaired *t* test with Welch’s correction (for Gaussian-distributed data) or an unpaired Mann-Whitney test (*, P<0.05; **, P<0.01; ***, P<0.001; ****, P<0.0001; ns, not significant). *R^2^* values indicate goodness-of-fit for the mathematical solutions. See also Figure S9 and Tables S2 and S3.

We previously showed that when the genetic dose of Plk4 was halved the centrioles grew slowly, but for a longer period of time, and so reached their normal size (Aydogan et al., 2018) – consistent with our observation that the Plk4-NG oscillator adapts in *Plk4-NG*^1/2^ embryos by reducing *A* and increasing *T* to maintain a relatively constant Ω. From the characteristics of the Plk4-NG oscillation in *asl^1/2^* embryos, we would predict that daughter centrioles should grow more slowly (as *A* is decreased), but for a normal period (as *T* is unchanged), and so centrioles would be too short (as *Ω* decreases). We measured the parameters of daughter centriole growth in *asl^1/2^* embryos and confirmed that this was indeed the case (Figure 6C). Together, these experiments support our hypothesis that the Plk4 oscillatory network helps to maintain a constant centriole size.

Importantly, these observations indicate that the threshold level of centriolar Plk4 required to promote centriole growth can vary depending on the conditions. In *asl^1/2^* embryos, for example, the Plk4 threshold that promotes centriole growth must be lower than in WT embryos – as centrioles grow for the same period as controls (although more slowly), even though less Plk4 is present at centrioles at any given time point. This is presumably because at the site of centriole assembly, the ratio of Asl to one or more of the other factors that promote centriole growth (such as Plk4, Ana2 or Sas-6) is altered in the *asl^1/2^* embryos. The lower levels of Plk4 at centrioles in *asl^1/2^* embryos could, for example, allow each Plk4 molecule to bind to Ana2 more efficiently (as there is less competition for Ana2) and so generate phosphorylated Ana2 more efficiently – thus lowering the threshold level of centriolar Plk4 required for centriole growth.

### Plk4 oscillations are detectable in non-dividing mouse liver cells and can be entrained by the circadian clock

In cyanobacteria and mammalian cells *in vivo*, the CCO can be entrained to the circadian clock (Matsuo et al., 2003; Yang et al., 2010). The Plk4 oscillator is normally entrained by the CCO, but it can autonomously promote centriole duplication even in the absence of a robust CCO (Figure 4). We wondered, therefore, whether the Plk4 oscillator could also be entrained by the circadian clock. To test this possibility, we examined a recently published diurnal proteome from non-regenerating mouse liver (Wang et al., 2018), where hepatocytes, the major building blocks of the hepatic tissue, are largely quiescent under normal conditions (Friedman, 2000). Several key cell cycle regulators (such as Cdk1, Cyclin E, Cyclin B1 and Plk1) were not detectable at any stage of the diurnal cycle, confirming that these cells were largely quiescent. In contrast, Plk4 was detectable and exhibited a clear oscillation that was entrained to the light/dark cycles (Figure 7). We presume that this level of Plk4 oscillation is sub-threshold for centriole biogenesis – as centrioles should not be duplicating in these non-dividing cells – but rather reflects Plk4’s ability to form an oscillator that runs independently of a robust CCO and that is instead entrained by the circadian clock.

**Figure 7.**
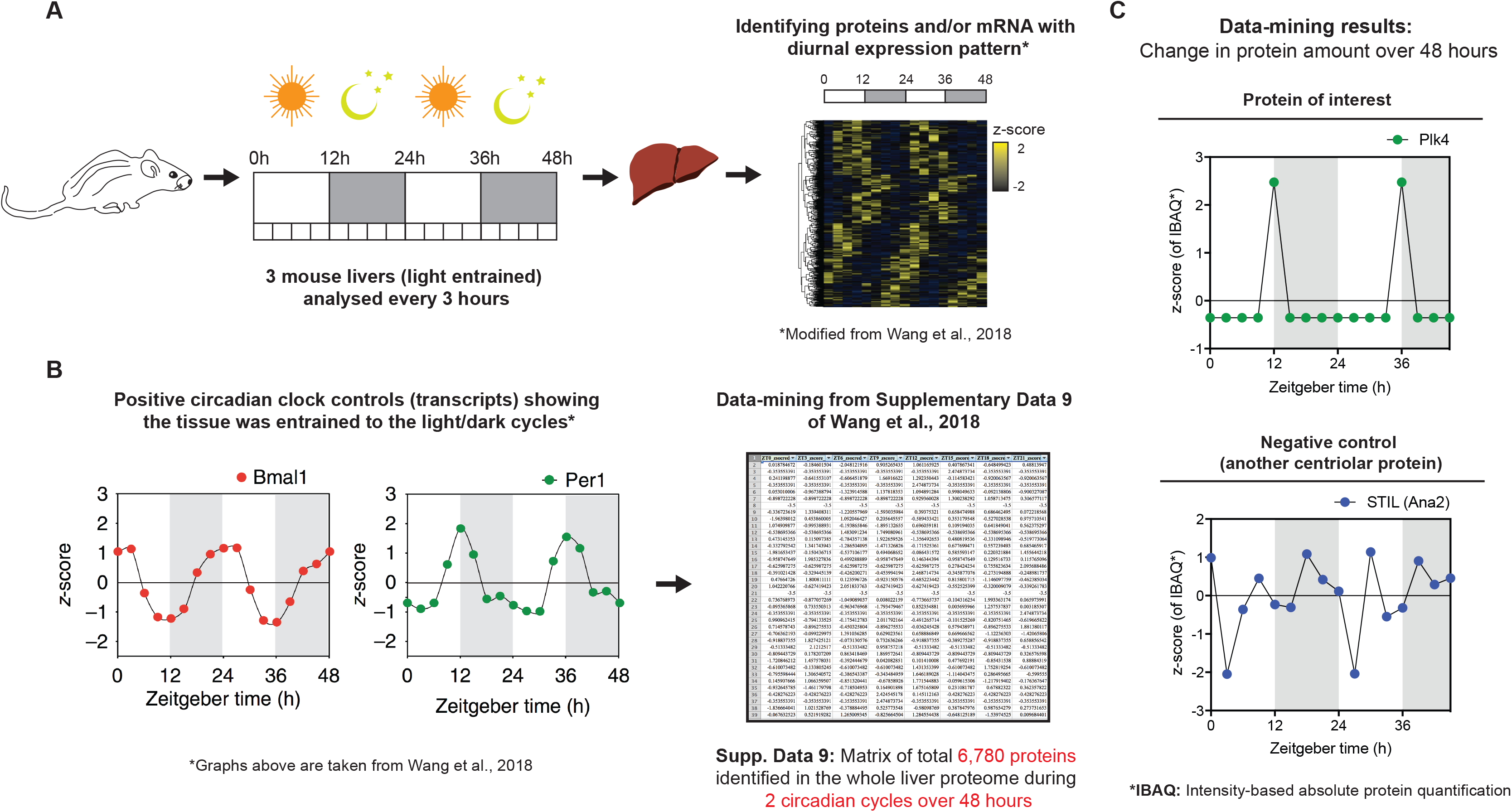
Plk4 may form an autonomous oscillator in mammals that can be entrained by the circadian clock. **(A)** Diagram shows the workflow used by Wang et al. to obtain a diurnal proteome of the whole liver of light/dark-entrained mice (Wang et al., 2018). **(B)** Graphs reproduced from Wang et al. (2018), show the relative diurnal expression of the circadian clock transcripts Bmal1 and Per1 as internal controls. We re-analysed the diurnal proteome produced in this study – comprising a matrix of z-scores for 6,780 proteins identified during 2 circadian cycles (Supplementary Data Set #9 from Wang et al., 2018). **(C)** Graphs we derived show the relative protein levels of Plk4 and the cartwheel component STIL (Ana2 in flies). Plk4 levels strongly spike in a periodic manner every circadian cycle, whereas STIL levels appear to randomly fluctuate and show neither a discernible pattern of oscillation nor any entrainment to the circadian clock. As these cells are generally not proliferating, centrioles should not be duplicating here, so the Plk4 oscillations are presumably sub-threshold for centriole biogenesis. We conclude that Plk4 levels naturally oscillate in quiescent liver cells, and the circadian clock entrains the period of these oscillations. Thus, Plk4 appears to form an autonomous oscillator in nondividing mammalian cells that is likely to be entrained by both the cell cycle and circadian clocks.

### Concluding remarks

Centrioles comprise two main compartments, an inner cartwheel structure and an outer set of microtubule (MT) blades (Guichard et al., 2013). In flies, the cartwheel and MT blades grow to similar sizes with broadly similar kinetics (Aydogan et al., 2018). In many other organisms, however, the MT blades extend past the cartwheel, and it is unclear how the growth of these two components of the centriole are normally coordinated to achieve a constant ratio between their ultimate sizes. The autonomous Plk4 oscillator regulates the growth of cartwheel structure, and we speculate that there must be a separate system that times and regulates the growth of MT blades. We suspect that this system must coordinate with the Plk4 oscillator to ensure that the cartwheel and MTs grow to the correct relative sizes. Identifying this second system, and understanding how it interacts with the Plk4 oscillator, may provide important clues as to how cells coordinate the growth of their often topologically-complex organelles.

There is great interest in determining the physical and molecular principles that cells use to regulate the biogenesis of their organelles (Liu et al., 2018; Mukherji and O’Shea, 2014). The idea that an organelle-specific autonomous oscillator could time and execute organelle biogenesis has, to our knowledge, not been proposed previously. We suggest that the autonomous Plk4 centriole oscillator could exemplify a more general mechanism for regulating organelle biogenesis, whereby autonomous oscillations in the levels/activity of key regulatory factors essential for organelle biogenesis could precisely time the initiation and duration of the growth process to ensure that organelles grow at the right time and to the appropriate size. Such oscillations would likely be phase-locked to either or both of the cell cycle and circadian clocks to precisely coordinate organelle biogenesis with other cellular events and processes.

## Abbreviations and symbols

*A*: Amplitude
*T*: Full width at half maximum
Ω: Area under the curve
CCO: Cdk-cyclin oscillator

## Acknowledgements

We are grateful to Laura Hankins, Fabio Echegaray, Drs. Marjorie Fournier, Christoffer Lagerholm, Pedro Carvalho, Bela Novak and Alain Goriely for advice and discussion, and Alain Goriely, Alissa Kleinnijenhuis and members of the Raff laboratory for critically reading the manuscript. Microscopy was performed at the Micron Oxford Advanced Bioimaging Unit, funded by a Strategic Award from the Wellcome Trust (107457). The research was funded by a Wellcome Trust Senior Investigator Award (104575; T.L. Steinacker, M. Mofatteh, A. Wainman, S. Saurya, J.W. Raff), an Edward Penley Abraham Scholarship (to M.G. Aydogan and L. Gartenmann), a BTG Junior Research Fellowship in Biomedical Sciences (Lincoln College, Oxford; to M. Mofatteh), a Cancer Research UK Oxford Centre Prize DPhil Studentship (C5255/A23225), a Balliol Jason Hu Scholarship, a Clarendon Scholarship (to S.S. Wong), Ludwig Institute for Cancer Research funding (to F.Y. Zhou), the St. Cross Emanoel Lee Junior Research Fellowship and a Biotechnology and Biological Sciences Research Council grant BB/N016858/1 (to M.A. Boemo).

## Author contributions

This study was conceptualised by M.G.A., M.A.B. and J.W.R. Investigation was done by M.G.A., T.L.S., M.M., L.G., A.W., S.S., and M.A.B. Data were analysed by M.G.A., T.L.S., L.G., F.Y.Z. and M.B.A. Methodology was developed by M.G.A., T.L.S., M.M., S.S.W., F.Y.Z., M.A.B. and J.W.R. Project was administered by M.G.A., M.A.B and J.W.R. Resources were shared/made by M.G.A., M.M., L.G., A.W., S.S., S.S.W. and M.A.B. Software development was carried out by M.G.A., T.L.S., S.S.W., F.Y.Z., and M.A.B. Overall supervision was done by M.G.A. and J.W.R. Validation experiments/analyses were carried out by M.G.A., A.W., S.S. and J.W.R. Finally, M.G.A., M.A.B. and J.W.R. wrote, reviewed and edited the draft.

## Materials and Methods

### *D. melanogaster* stocks and husbandry

The specific *D. melanogaster* stocks used in this study are listed in Table S4, and the lines generated and tested here are listed in Table S5. To generate Plk4-mNeonGreen and Asl-mKate2 constructs: **1)** NheI restriction enzyme sites were introduced into an mCherry C-terminal Gateway vector (Basto et al., 2008), using the Quikchange II XL mutagenesis kit (Agilent Technologies). **2)** The mCherry tag was replaced with either mNeonGreen (Shaner et al., 2013) (Allele Biotechnology) or mKate2 (Shcherbo et al., 2009) tags by homologous recombination via In-fusion Cloning (TaKaRa). **3)** NheI restriction enzyme sites were removed via site-directed mutagenesis, using the Quikchange II XL mutagenesis kit (Agilent Technologies). These vectors were recombined via Gateway technology to pDONR-Zeo vectors (Thermo Fisher Scientific) where the genetic regions of either *Plk4* (Aydogan et al., 2018) or *asl* (Novak et al., 2014) were previously cloned from 2 kb upstream of the start codon up to (but excluding) the stop codon. Primer sequences are listed in Table S6. Transgenic lines were generated using standard P-element mediated transformation by the Fly Facility in the Department of Genetics, University of Cambridge (Cambridge, England, UK). Flies were maintained at 18°C or 25°C on *Drosophila* culture medium (0.77% agar, 6.9% maize, 0.8% soya, 1.4% yeast, 6.9% malt, 1.9% molasses, 0.5% propionic acid, 0.03% ortho-phosphoric acid, and 0.3% nipagin) in vials or bottles.

### Hatching experiments

To measure embryo hatching rates, 0-3 h embryos were collected and aged for 24 h, and the % of embryos that hatched out of their chorion was calculated.

### Synthesis of double-stranded RNA

Double-stranded RNAs (dsRNAs) against cyclins A, B and B3 were synthesised essentially as described previously (McCleland and O’Farrell, 2008). Primer sequences used for gene amplification are listed in Table S6. The linker sequence 5’-GGGCGGGT-3’ was added to the 5’ end of each primer to facilitate the addition of the T7 promoter in the second round of PCR using a universal primer containing the linker (5’-TAATACGACTCACTATAGGGAGACCACGGGCGGGT-3’). The resulting RNA was precipitated with 8 μl of 3M Na-Acetate and 220 μl of 100% ethanol before washing with 70% cold ethanol. The RNA pellets were air dried and resuspended in 30 μl of RNase-free diethylpyrocarbonate-treated water (Thermo Fisher Scientific). To generate double-stranded molecules, RNAs were placed in a 67.5°C water bath for 30 min, and allowed to cool to room temperature over 90 min. Unincorporated UTPs were removed using CHROMA SPIN-100-DEPC-H_2_O columns (Clontech) according to the manufacturer’s instructions. To confirm the synthesis of the correct RNA product, 3 μl of the final reaction was subjected to electrophoresis on a 1.5% agarose gel using 2xRNA loading buffer (Thermo Fisher Scientific). A 1:1 mix of RNA and loading buffer was heated to 65°C for 5 min and then placed on ice to denature any secondary structure of RNA.

### Embryo collections and drug and dsRNA injections

For embryo collections, 25% cranberry-raspberry juice plates (2% sucrose and 1.8% agar with a drop of yeast suspension) were used. Embryos for imaging experiments were collected for 1h at 25°C, and aged at 25°C for ~45–60 min. Embryos were dechorionated by hand, mounted on a strip of glue on a 35-mm glass-bottom Petri dish with 14 mm micro-well (MatTek), and were left to desiccate for 1 min at 25°C. After desiccation, the embryos were covered with Voltalef grade H10S oil (Arkema). Embryos for drug or dsRNA injection experiments were treated in the same way except that the desiccation period was increased to 5-6 min. Embryos were injected with dsRNA at a needle concentration of 0.6–0.8 mg/ml; colchicine in Schneider’s insect medium was injected at a needle concentration of 100 μg/ml (diluted from a 20 mg/ml stock in water) (Raff and Glover, 1989).

### Immunoblotting

Immunoblotting was performed as described previously (Aydogan et al., 2018). Primary antibodies used in this study are as follows: mouse anti-GFP (Roche) and mouse anti-Actin (Sigma). Both the antibodies were used at 1:500 dilution in blocking solution (Aydogan et al., 2018). For all blots, 10, 20 or 30 staged early embryos were boiled in sample buffer and loaded in each lane. The incubation period for primary antibodies was 1 h (or overnight at 4°C). Membranes were quickly washed 3x in TBST (TBS and 0.1% Tween 20) and then incubated with HRPO-linked anti-mouse IgG (both GE Healthcare) diluted 1:3,000 in blocking solution for 45 min. Membranes were washed 3×15min in TBST and then incubated in SuperSignal West Femto Maximum Sensitivity Substrate (Thermo Fisher Scientific). Membranes were exposed to film using exposure times that ranged from <1 to 60s.

### Image acquisition, processing, and analysis

#### Spinning disk confocal microscopy

Living embryos were imaged at room temperature using a system equipped with an EM-CCD Andor iXon+ camera on a Nikon Eclipse TE200-E microscope using a Plan-Apochromat 60x/1.42-NA oil DIC lens, controlled with Andor IQ2 software. Confocal sections of 17 slices at 0.5μm intervals were collected every 30 s. A 488nm laser was used to excite mNeonGreen and GFP, and a 568nm laser was used to excite mCherry. Emission discrimination filters were applied when mNeonGreen and mCherry were imaged together.

Post-acquisition image processing was performed using Fiji (National Institutes of Health). Maximum-intensity projections of the images were first bleach-corrected with Fiji’s *exponential fit* algorithm, and background was subtracted using the *subtract background* tool with a rolling ball radius of 10 pixels. Plk4-NG, Sas-6-mCherry or -GFP, and Asl-mCherry or -GFP were tracked using the Fiji plug-in TrackMate (Tinevez et al., 2016) with a track spot diameter size of 1.1 μm. The regressions for the centriole growth curves (Sas-6-GFP or -mCherry) were calculated in Prism 7 (GraphPad Software), as described previously (Aydogan et al., 2018). The regressions for the Plk4 oscillation curves (Plk4-NG) were calculated using the *nonlinear regression* (curve fit) function in Prism 7. Discrete Plk4 oscillation curves in S-phase were initially fitted against four different functions to assess the most suitable regression model: 1) Lorentzian, 2) Gaussian, 3) Increase – Constant – Decrease, and 4) Increase – Decrease. Among these models, Lorentzian best fit the data (Figure S1D), so all the discrete Plk4 oscillation curves in S-phase were regressed using this function. The Lorentzian and Gaussian functions are described in Prism 7, while the latter two functions are in-house algorithms (Alvarez Rodrigo et al., 2018).

In order to plot the dynamics of Plk4-NG and Sas-6-mCherry together (Figures 2, S3A and S3B), the highest mean fluorescence signal for each tag was normalized to 1 and was accordingly scaled across cycles 11-13 (the scaling factor for Plk4-NG was calculated from the data shown in Figures 1B and 1C). Note that the amplitude of the Plk4 oscillation does not appear to decrease significantly between nuclear cycles 11-13 in the data shown in Figure 2A—in contrast to the Plk4 oscillations shown in Figure 1B. This is not because of the scaling procedure applied to the data shown in Figure 2A (described above), but rather because embryos that failed to grow their centrioles were excluded from the analysis shown in Figure 2A. The amplitude of the Plk4 oscillations was lower in these embryos (Figure 2C), so embryos with low amplitude Plk4 oscillations were effectively excluded from the analysis shown in Figure 2A. Almost all of these excluded embryos were at nuclear cycle 13, so the “average” oscillation at nuclear cycle 13 is in reality an average of only those embryos that had a relatively high amplitude Plk4 oscillation.

In all the imaging experiments, the beginning of S-phase was taken as the time at which the old and new mother centrioles were first detected to separate from each other (termed “centrosome separation” or “CS”). Entry into mitosis was taken as the time of nuclear envelope breakdown (NEB), which could be determined in our movies by adjusting the contrast to visualise when the cytoplasmic pool of the fluorescent protein was first observed to enter into the nucleus.

#### Analysis of centriole “fertility” in embryos injected with cyclin A-B-B3 dsRNA

In experiments where we depleted embryos of mitotic cyclins during early rounds of nuclear division, we observed qualitatively that “fertile” centrioles exhibited distinct Plk4-NG fluorescence peaks that often appeared to correlate with centriole duplication events, while “sterile” centrioles, which did not duplicate during the imaging period, exhibited no obvious peaks (Figure 4B). To test if we could more quantitatively distinguish between fertile and sterile centrioles, we analysed all 81 centrioles that we could track throughout the observation period in 3 different embryos. We first assessed the average signal-to-noise ratio (SNR) of Plk4-NG fluorescence of each centriole over the entire observation period and found that fertile centrioles exhibited a significantly higher SNR than sterile centrioles—assessed using a t-test assuming equal variance (Figure S4B). The distribution of SNR within sterile and fertile centriole signals was unimodal and symmetrically distributed (Figure S4C), so we attempted to classify centrioles in an unbiased way by thresholding the SNR. Based on the bimodality of the SNR, an automatic threshold was determined from the data using Otsu thresholding (*red dashed* line Figure S4C); the classification performance was summarized in a visual confusion matrix, which shows the proportion of correctly and falsely classified signals (Figure S4D). This unbiased computational method successfully classified ~74% of the fertile centrioles and ~71% of the sterile centrioles.

We next tested whether computationally identified peaks in the Plk4-NG signal were correlated with centriole duplication events. Peaks or local maxima in the Plk4-NG fluorescence signals were initially detected as points whose two direct neighbouring points have lower amplitude. This was implemented using Scipy’s *find_peaks* function (Jones et al., 2001) with parameters, height=0, distance=1, prominence=0.1 (minimum). Direct application to the raw signal identified too many false positives, so the raw signal was first processed using an asymmetric least squares filter (Eilers and Boelens, 2005) with parameters, λ=10^2^, p=0.1 run for 5 iterations—see Figure 4B for examples of how the raw peak data (*black line*) is converted to the filtered peak data (*red line*) by this process. To determine whether the filtered Plk4-NG peaks were predictive of centriole duplication, we determined all the peaks for the fertile centriole signals and assessed whether these peaks could be used to “retrieve” the real or relevant time points for centriole duplication. The performance of such retrieval can be evaluated using “precision” (the number of relevant retrievals amongst all retrieved instances—in this case the number of Plk4 peaks associated with a centriole duplication event divided by the total number of Plk4 peaks) and “recall” (the number of relevant instances retrieved of the total relevant instances—in this case the number of Plk4 peaks associated with a centriole duplication event divided by the total number of centriole duplication events), as defined below.

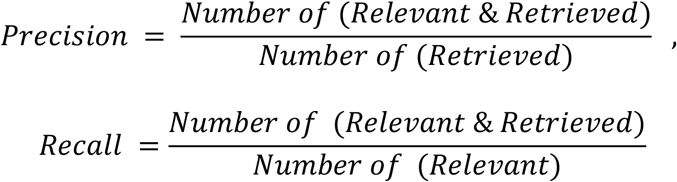

The evaluation of such a system naturally depends on the cut-off to call a positive match between the Plk4-NG signal peak and its corresponding centriole duplication time. Too small a cut-off (e.g. 0 minutes) is unrealistic: no system can predict time perfectly; while too large a cut-off (e.g. 15 min) is too lenient and non-specific. Figure 4C plots the precision evaluated over all centrioles for different temporal cut-offs attempting to uniquely match Plk4-NG peaks to the nearest duplication time within a given time window. The elbow point (*red dashed* line) at 5 min was selected as an appropriate cut-off with a precision of ~80%. (Note that the recall was not plotted in this graph, but it exhibits a similar behaviour to the precision: the *Number of (Relevant)* = 52, while the *Number of (Retrieved)* = 49). This temporal cut-off can also be interpreted as an estimate of the temporal accuracy to which Plk4-NG peak time associates with centriole duplication time. For comparison, we also derived the mean temporal separation distance of peaks and duplication events, if the same number of experimental centriole duplication times were randomly distributed over the same time interval for each embryo. 1000 simulations were run per embryo to produce a distribution. Across all embryos, an average temporal separation distance (for randomly distributed duplication times) was 10.5 minutes (data not shown), twice as long as the chosen 5 min cut-off, thus the association is not coincidental.

In addition, we assessed the precision and recall performance over different possible threshold values (based on peak prominence) used to call a Plk4 ‘peak’. To do this, we computed the precision-recall curve. All detected Plk4 peaks (*Black* dots; Figure 4E) were ranked according to their peak prominence from high to low and were assigned uniquely to a duplication event according to a 5 min time window for determining a positive match. Peaks that could not be uniquely assigned in such a manner were regarded as ‘negatives’. The graph then plots the precision, recall values if the threshold for calling a peak were set as the peak prominence value of each peak in descending order. Beyond the detected peak associated with a peak prominence of 0.12 (i.e. points right of this point), the precision drops sharply. At this threshold, precision and recall are jointly optimised. This suggests that a minimum level of Plk4-NG peak fluorescence intensity is required to predict duplication. The ability of Plk4 peaks to predict duplication across all peak prominences (over the selected time window of 5 min) is quantified by the integrated area under the curve or average precision (AP_5min_). If there were no overall correlation between a Plk4 peak and a duplication event, AP_5min_ would be 51.5% (given by # duplications / (# duplications + # peaks)); the score of ~75% indicates a strong overall correlation (Figure 4E).

Finally, the correlation between the Plk4-NG peaks and times of centriole duplication was examined (Figure 4F), which provided an alternative accuracy test. Plk4-NG peaks were uniquely matched to the nearest centriole division times without using a temporal cut-off over individual centrioles from three independent embryos. Pearson correlation r, r-squared R^2^ and *P* values are reported as goodness of fit. The fitted regression line, *y* = 0.87*x* + 3.69. Together, these unbiased computational analyses indicate that the Plk4 oscillations at individual centrioles are highly correlated with the time at which these centrioles duplicate.

#### 3D-Structured Illumination Microscopy (3D-SIM)

Living embryos were imaged at room temperature using a DeltaVision OMX V3 Blaze microscope (GE Healthcare). The system was equipped with a 60x/1.42-NA oil UPlanSApo objective (Olympus Corp.), 488nm and 593nm diode lasers, and Edge 5.5 sCMOS cameras (PCO). Spherical aberration was reduced by matching the refractive index of the immersion oil (1.514) to that of the embryos. 3D-SIM image stacks consisting of six slices at 0.125μm intervals were acquired in five phases and from three angles per slice. The raw acquisition was reconstructed using softWoRx 6.1 (GE Healthcare) with a Wiener filter setting of 0.006 and channel-specific optical transfer functions (OTFs). Filters used for the green and red channels were a 540/80 centre band pass filter and a 605-long pass filter, respectively. For two-colour 3D-SIM, images from green and red channels were registered with the alignment coordination information obtained from the calibrations using 0.2μm-diameter TetraSpeck beads (Thermo Fisher Scientific) in the OMX Editor software. The SIMCheck plug-in in ImageJ (National Institutes of Health) was used to assess the quality of the SIM reconstructions (Ball et al., 2015); only images that passed this test were used.

### Mathematical modelling and its experimental validation

#### The mathematical model

Figures 5A and 5B, specify a regulatory network wherein Plk4 binds to an Asl receptor with high affinity; this activates Plk4, allowing it to phosphorylate itself and Asl multiple times. After a certain number of phosphorylations, Asl switches to a new state that binds Plk4 with low affinity. As a result, Plk4 unbinds, leaving Asl in a phosphorylated, low affinity state. In our model (which only attempts to model the Plk4 oscillation through a single cycle of S-phase), Asl is not dephosphorylated after Plk4 dissociates. We suspect that, in reality, Asl is normally dephosphorylated in early mitosis, which “resets” it to a high-affinity state in preparation for the next oscillation. In this model, the gradual conversion of centriolar Asl to a low affinity binding state forms a time-delayed negative feedback loop, wherein Asl effectively activates Plk4 to gradually promote its own inhibition. After making assumptions about the chemical kinetics of the system and imposing suitable initial conditions, the behavior of this regulatory network can be simulated by mapping it onto a set of coupled ordinary differential equations (see below).

In the model, it is assumed that Plk4 is well-mixed in the cytoplasm and that centrioles are large macromolecular structures; this allows Plk4 and Asl receptors on the centriole to follow mass-action kinetics. [***P***] and [***A***] denote the concentrations of unbound cytoplasmic Plk4 and unbound centriolar Asl, respectively. Plk4 binds to Asl with the fixed rate constant ***k***, and the rate constant of the reverse reaction is sufficiently small that any unbinding is ignored. Once Plk4 is bound to Asl, it can only unbind after it has phosphorylated Asl nine times (see below for an explanation of why this number was chosen). [***A*_0_**] denotes the concentration of Asl receptor that has bound Plk4, but has not yet been phosphorylated, while [***A_i_***] denotes the concentration of Asl receptors that have been phosphorylated *i* times. Each phosphorylation of Asl by Asl-bound Plk4 has rate constant ***k***_2_ and, following nine phosphorylations, the Asl is switched to a state that binds Plk4 with very low affinity [***A*_9_**]. Once Asl has been converted to this low affinity state, Plk4 unbinds at rate ***k*_3_**. The rate constant of the reaction where Plk4 binds to Asl receptor in this low affinity state is assumed to be sufficiently small that this reaction is ignored in the model.

Intuitively, *k* scales the affinity with which Plk4 binds to an Asl receptor. By mass action kinetics, the rate of this reaction is given by *k* * [*P*] * [*A*]. It is assumed in the model that Plk4 is abundant enough in the cytoplasm that its concentration does not decrease over the few minutes of the single S-phase cycle; this assumption means that [*P*] is constant (see below for a further explanation of why we believe this assumption is valid). Therefore, the number of parameters in the model is reduced by introducing the new rate constant ***k*_1_** = ***k*** * [***P***].

Using the assumptions above, the regulatory network in Fig. 4, A and B, is simulated using the following set of ordinary differential equations which are solved over the time domain 0 ≤ t ≤ S, where S is the length of S-phase:

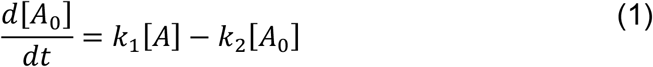

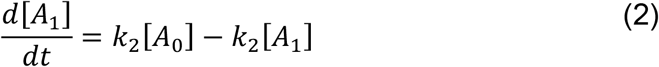

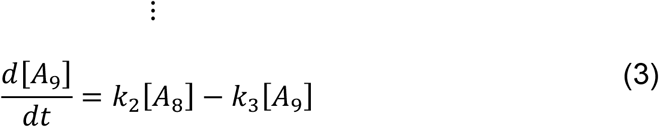

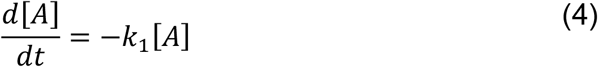

Appropriate initial conditions at t = 0 are,

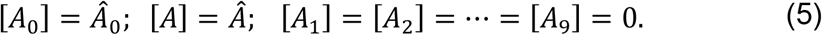

In equation (5), the positive constant ***Â*_0_** is the initial amount of Plk4-bound Asl at the centriole at the start of each S-phase, which is determined experimentally for each cell cycle using the techniques described in the *Image acquisition, processing and analysis* section of the *Materials and Methods*. The constant ***Â*** is the initial amount of unbound Asl, so the total amount of Asl in the system is given by *A_tot_* = *Â*_0_ + *Â*.

The model specified by equations (1)–(5) has an analytical solution, which is solved using MATLAB’s *dsolve*. Values for the parameters *Â, k*_1_, *k*_2_, and *k*_3_ were determined by fitting the curve [*A*_0_](*t*) + [*A*_1_](*t*) + ··· + [*A*_9_](*t*) to the experimentally measured data for the amount of Asl-bound Plk4 (i.e., the Plk4 that is recruited to the centriole) over time. Fitting was done using a *trust-region* algorithm to optimise a nonlinear least squares penalty function.

The fitting was constrained so that all parameters were positive, and *k*_1_, *k*_2_, and *k*_3_ were between 0 and 1. Parameter values are shown in Tables S1–S3. As explained below, the solutions to this model are very insensitive to variations in *k*_3_ (see Figure S6), so in the solutions presented here *k*_3_ was kept at a constant value of 0.06906, which was the best-fit parameter value for cycle 12 (Tables S1–S3).

In the model, we assumed that Asl had to be phosphorylated by Plk4 nine times before it switched to a low-affinity state—indicated by variables [*A*_0_], ···, [*A*_9_]. We chose this number because we tested the effect of the number of phosphorylation sites on the model solution. The best fit curves for [*A*_0_](*t*) + ··· + [*A*_1_](*t*) + ··· + [*A_N_*](*t*) for N = 1, 4, 9, 14, or 16 suggested that the model fits the data slightly better when setting N=9 (*R*^2^=0.9996) as opposed to N=1 (*R*^2^=0.9152), N=4 (*R*^2^=0.9886), N=14 (*R*^2^=0.9962) or N=16 (*R*^2^=0.9931), so we used 9 phosphorylation sites in all subsequent modelling. Interestingly, however, the curves for N=4, 9 or 14 were all within the standard error of the mean, indicating that the model is relatively insensitive to the choice of the number of phosphorylation sites within this range (data not shown).

Table S1 shows that the *trust-region* algorithm finds a good fit (*R*^2^>0.99) for the model to the experimental data (Figure 5C), but this provides little information about uniqueness of the fit: There may be other subsets of the parameter space that also provide a good fit to the data. To see if any such regions could be detected, the parameter space was further explored by using a *Metropolis-Hastings Markov chain Monte Carlo* algorithm. Four Markov chains were started at the positions in the parameter space specified in Table S7. Figure S6 shows the six two-dimensional traces of the four-dimensional parameter space. For clarity, only points that provided a good fit to the cycle 12 data (*R*^2^>0.95) are shown.

The results in Figure S6 reveal how sensitive the model is to changes in each parameter value. Parameter *k*_3_ is very insensitive, so we set it to a constant arbitrary value of 0.06906 to allow a more intuitive comparison of the other variables in Table S1–S3. In contrast, the model only fit the data well for a relatively narrow range of values for *k*_1_, *k*_2_, and *Â*. The Monte Carlo results also reveal correlations between *k*_1_ and *k*_2_, *k*_1_ and *Â*, and *k*_2_ and *Â*. For example, these results show that if *Â* (the initial amount of unbound Asl receptor at the start of S-phase) is reduced, the model can still fit the data well if *k*_2_ is decreased and *k*_1_ is increased. While these results suggest that there is a single, continuous region of the parameter space that provides a good fit to the data, it is still possible that there are other such regions that the Markov chains in Figure S6 did not explore. However, the results (in Figure S6B) show that the points, which are identified at the center of the parameter region, provide the best fit to the data. This suggests the nonlinear least squares minima found by the *trust-region* fitting is unlikely to be very sensitive to the initial seed that was chosen.

All the equations used for mathematical modelling and regressions are available in the following web link: <https://github.com/MBoemo/centriole_oscillator_model.git>. See below for further information about how this model was used to probe the effect of reducing the concentration of Plk4 and Asl, shown in Tables S2 and S3.

#### Experimental validation of the mathematical model

To assess the validity of our mathematical model (Figures 5A and 5B), the model was first used to individually fit the discrete Plk4-NG oscillation data from S-phase of cycles 11, 12 and 13 (Figure 5C). The best-fit parameter values fit the data for each cycle well (*R*^2^≥0.99) and generated biologically plausible parameters for *k*_1_, *k*_2_ and *k*_3_, and the total amount of Asl receptor at the centriole (*A_tot_*) (Table S1). Note that *k*_2_ describes the rate at which Plk4 phosphorylates Asl: as a simplification, it was assumed that *k*_2_ is constant for each of the multiple phosphorylation sites in Asl, and that the bound Plk4 molecule sequentially phosphorylates one site after another, with no dephosphorylation possible (as we speculate that the dephosphorylation of Asl is regulated and only occurs at the start of mitosis, see *Results and Discussion*). The parameter space was probed for any parameter values that would provide a good fit to the data by using Markov chain Monte Carlo methods. For all parameters except *k*_3_, the best-fit values occupied a single, narrow region of the parameter space. The Monte Carlo results revealed that parameter *k*_3_ was very insensitive and could provide a good fit to the data for a broad range of values (Figure S6). This is presumably because the rate of phosphorylating Asl at multiple sites is relatively slow compared to the rate at which Plk4 is subsequently released from the multiply phosphorylated Asl—so the rate of release is not limiting. To ease the comparison between different parameters, *k*_3_ was therefore held at an arbitrary constant value in all the models shown here (Tables S1–S3).

Interestingly, the best-fit parameters for cycles 11–13 showed that the biggest difference between the parameters of the Plk4 oscillations at each cycle is in *k*_1_—the rate at which Plk4 binds to Asl (which is dependent on the cytoplasmic concentration of Plk4). Although our model assumes that the cytoplasmic concentration of Plk4 remains constant during the S-phase period within each cycle, if the phosphorylated Plk4 molecules that are released from the Asl receptor are ultimately degraded—and there is good evidence that Asl activates Plk4 to promote Plk4 degradation (Klebba et al., 2015)—there could be a wave of phosphorylated-Plk4 degradation in the cytoplasm towards the end of S-phase. If so, the cytoplasmic levels of Plk4 would get successively lower at the start of each successive cycle, as our PeCoS analysis indicates is the case (Figure 5F).

The effects of reducing the genetic dose of *Plk4* by half (*Plk4-NG^1/2^* embryos—see *Results and Discussion* for details) were analysed. Our PeCoS analysis indicated that there was a ~45% drop in the cytosolic levels of the Plk4-NG protein in the *Plk4-NG^1/2^* embryos. When the model was fit to *Plk4-NG^1/2^* oscillation, the best-fit (*R*^2^=0.996) parameter had a *k*_1_ value that was ~39% of the control value (Table S2), so in reasonable agreement with our experimental measurements. These parameter values also suggested that the total amount of centriolar Asl (*A_tot_*) should remain relatively unchanged between the *Plk4-NG* and *Plk4-NG^12^* conditions (Table S2). Centriolar Asl levels were analysed in embryos expressing Asl-mCherry in either WT vs *Plk4^1/2^* conditions, and our findings showed that this was indeed the case (Figure S9A).

Next, the effects of reducing the genetic dose of *asl* by half (*asl^1/2^* embryos— see *Results and Discussion* for details) were analysed. Interestingly, the best-fit parameter values (*R*^2^=0.999) predicted that the total centriolar Asl levels (A_tot_) would be reduced by only ~28% in *asl^1/2^* embryos (Table S3). This value was therefore directly measured in embryos expressing either one or two copies of Asl-GFP (under the control of its own promoter in an *asl* mutant background), and encouragingly our findings showed that reducing the genetic dose of Asl-GFP by half led to a reduction of only ~30-35% in centriolar Asl-GFP levels (Figure S9B). Moreover, the parameter values suggested that the concentration of Plk4 (incorporated in the *k*_1_ term) should not vary significantly between WT and *asl^1/2^* conditions (Table S3). We tested this prediction of the model by blotting for Plk4-GFP (transgenically expressed from its own promoter in a *Plk4* mutant background) in control and *asl^1/2^* embryos. This revealed that the amount of Plk4-GFP did not change dramatically between the control and *asl^1/2^* conditions (Figure S9C).

Taken together, these analyses indicate that our model can robustly describe the Plk4-NG oscillations under normal conditions (Figure 5C) and when the levels of either Plk4 or Asl are perturbed experimentally (Figures 6A and 6B). Moreover, the model makes several plausible predictions about the relative levels of these proteins in the perturbed conditions that are close to the levels that were measured experimentally.

Finally, the best-fit value of *k*_2_ (reflecting the kinase activity of individual Plk4 molecules) decreased slightly between cycles 11 to 12 and decreased more significantly between cycles 12 to 13 (by ~9% and ~37%, respectively); *k*_2_ also decreased when levels of Plk4 were genetically reduced in *Plk4-NG^1/2^* embryos (by ~25%)—but not when Asl levels were genetically reduced in *asl^1/2^* embryos (Tables S1–S3). The molecular basis for this inferred decrease in kinase activity remains unknown, but we previously suggested that centriolar Plk4 might integrate several inputs at the start of each cycle (from, for example, cell cycle regulators, or its activator Ana2/STIL) and adjust its kinase activity in response to the lengthening of S-phase during successive nuclear cycles (Aydogan et al., 2018).

### Fluorescence Correlation Spectroscopy (FCS)

#### FCS setup and measurements

Point FCS measurements were performed on a confocal Zeiss LSM 880 (Argon laser excitation at 488 nm and GaASP detector) with the Zen Black Software. A C-Apochromat 40x/1.2 W objective and a pinhole setting of 1AU were used. A laser power of 10 μW was used, and no photobleaching was observed during the measurements. The microscope was kept at 25°C using the Zeiss inbuilt heating insert P and the heating unit XL. A schematic overview of the methodology used is shown in Figure S7A, and a comparison of the average autocorrelation curves generated at the start of S-phase of nuclear cycles 11-14 is shown in Figure S7B.

The effective volume of the imaging setup was estimated to be 0.28 fL by averaging the estimate obtained by three independent methods, as described previously (Rüttinger et al., 2008): **1)** Measuring the concentration of a soluble Alexa Fluor^™^ 488 NHS Ester dilution series (100 nM, 10 nM, 1 nM and 0.1 nM); **2)** Measuring the diffusion time for Alexa Fluor^™^ 488 NHS Ester (same concentrations) in water at 25 °C. The measured diffusion time was then compared to a previously reported diffusion coefficient for the Alexa Fluor^™^ 488 NHS Ester (Petrášek and Schwille, 2008); **3)** Imaging subresolution beads (FluoSpheres^™^ Carboxylate-Modified Microspheres, 0.1 μm) and determining the effective volume via Gaussian fitting with the *line tool* and *Z-axis profile* in ImageJ (Bethasda, USA).

Embryo collections (from mother flies expressing Asl-GFP under the control of its own promoter in an *asl* mutant background) were as described above, with the exception of using high precision 35 mm, high Glass Bottom μ-dishes (ibidi). Before every measurement, spherical aberrations were adjusted on the correction collar of the objective by maximising the count-rate per molecule (CPM). At the beginning of S-phase in each cell cycle (when the old and new mother centrioles were separating), consecutive cytoplasmic measurements were made 6x for 10 sec each at the centriolar plane of the embryo. Individual recordings where centrioles moved through the measurement spot, based on the highly erratic shape of the correlation curve (<2% of all recordings), were discarded.

#### Autocorrelation analysis and post-acquisition curve fitting

The autocorrelation function, *G*(*τ*), was calculated during each measurement in the Zen Black software using the following equation:

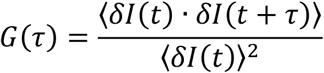

where *〈〉* denotes a time average, *δI*(*t*) describes the intensity fluctuation at the time point *t*, and *τ* states the lag time of the autocorrelation.

All 10 sec-recordings were then fitted with 8 different 3D diffusion models using the software FoCuS-Point (Waithe et al., 2016) with the following equation:

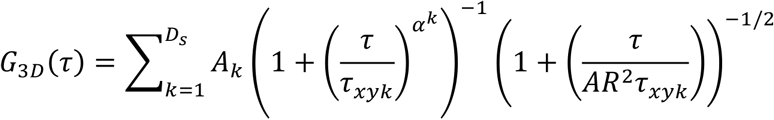

where *A_k_* defines the fraction of a diffusing species for which the sum of all diffusing species equals 1, *τ_xy_* describes the average residence time of the diffusing species in *V_eff_, α* accounts for anomalous subdiffusion within the cytoplasm, and *AR* is a structural parameter that describes the relationship among the x, y and z-axes of the excitation volume.

Dark states of the fluorophore were fitted with the following formula:

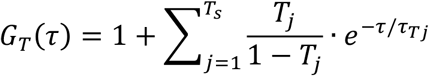

where *T* depicts the triplet population, and *τ_T_* states the triplet correlation time during which the fluorophore stays in the dark state (Schönle et al., 2014).

The data was fitted within the boundaries of 4×10^-4^ ms and 1.5×10^3^ ms, and the dark states were restricted to 10-300 μs for the blinking state, and 1-10 μs for the triplet state. The models (**Ms**) were defined as the following: **M1)** 1 diffusing species (ds) 0 blinking states (bs) 0 triplet states (ts); **M2)** 1ds 1bs 0ts; **M3)** 1ds 0bs 1ts; **M4)** 1ds 1bs 1ts; **M5)** 2ds 0bs 0ts; **M6)** 2ds 1bs 0ts; **M7)** 2ds 0bs 1ts; **M8)** 2ds 1bs 1ts. In all models, the structural parameter *AR* and the anomalous subdiffusion parameter *α* were kept constant at 5 and 0.7, respectively.

In order to avoid over-fitting the data, the most plausible model to describe the autocorrelation functions was selected using the Bayesian Information Criterion (BIC), which is based on the likelihood function, but introduces a penalty term for the complexity (number of variables) for the models (Schwarz, 1978). In this study, **M4** was the preferred model to describe Asl-GFP diffusion (Figure S7A_(iv)_). The concentration was calculated from the FoCuS-point fit data of the preferred model:

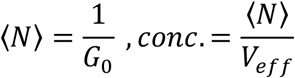

where *N* states the average number of particles within the effective volume *V_eff_*, and *G*_0_ represents the height of the autocorrelation function at t=0.

#### FCS background corrections

In order to estimate the contribution of the background noise, 22 wild-type embryos were measured with the same laser intensity (10 μW) and in roughly the same plane and developmental stage as the Asl-GFP embryos. Despite no observable correlated background, the uncorrelated background contributed ~30% of the total photon count rate, presumably due to the low concentration of cytosolic Asl-GFP and the high autofluorescence of the embryo itself (Figure S7_(vi)_). Background corrections were performed after the autocorrelation analysis by calculating the correction factor χ^2^ using the following formula (Koppel, 1974):

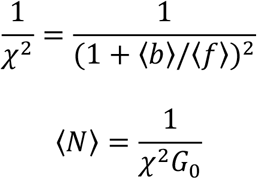

where 〈*b*〉 denotes the average background and 〈*f*〉 states the average count rate of the sample.

#### Data restriction

In some FCS measurements a sudden drop in CPM was observed, possibly due to movements within the embryo or the embryo drifting away from the measurement plane. When this happened, a strong, often unreasonable increase in concentration was observed. These outliers were therefore discarded based on a ROUT outlier test (with the aggression factor Q=1%), which was performed on all 10 sec-long concentration measurements (the *red* data points in Figure S7_(vi)_). Only the embryos with at least 4×10sec recordings (after discarding outliers and erratically shaped ACFs) were included in the final analysis.

### Peak Counting Spectroscopy (PeCoS)

FCS was not sensitive enough to investigate the cytosolic concentration of Plk4-NG as its concentration was too low. We therefore developed a new method that we term *Peak Counting Spectroscopy* (PeCoS) that allows the concentration of low abundance proteins to be measured accurately (Figure S8A). PeCoS uses the same set up as the point FCS protocol described above, but it differs in terms of its data acquisition and analysis. In PeCoS, the intensity peaks, which are generated by a fluorophore moving through the effective volume, are counted as a proxy for concentration. Due to the low cytosolic concentration of the fluorescently-tagged protein of interest (e.g., Plk4-NG in this study), spherical aberrations could not be corrected within the same embryo where the measurements were taken. Therefore, embryos that express a bright fluorescent centriolar marker were positioned next to the experimental embryos on the same imaging dish and these were used for correction collar adjustment (Figure S8A_(i)_). Experimental recordings were then captured for 180 sec (instead of 6×10 sec), as the number of particles that pass through the field of view was usually very low. Before every measurement, the observation region was pre-bleached with the same laser intensity (10 μW) for 3 sec to bleach away any potential immobile fraction.

Instead of autocorrelation analysis, the resulting intensity traces (Figure S8A_(iii)_) were quantified for their number of peaks, which originate from a fluorophore moving through the excitation volume and causing a detectable burst of photons. In order to determine the cut-off threshold, which was used to subtract the background noise, 40 control embryos were measured. These control embryos were from mothers expressing Asl-mKate to allow measurements at the centriolar plane and at the right nuclear cycle stage (beginning of S-phase). “Mean + n*SD” (where n = 1,2,3,…) of all control recordings was subtracted from each control recording, and the threshold that resulted in an average of less than five peaks (per 180 sec control measurement) was subtracted from all intensity traces (Figure S8A_(iv-vi)_). This threshold was found to be a good compromise for minimizing the background noise without discarding too much information. A Python script was written in order to automate this procedure, which is available via <https://github.com/SiuShingWong/PeCoS>. The subtraction of “Mean + 8*SD” resulted in an average peak count of 3.25 for the control recordings, and this was used for the background subtraction in all *in vivo* measurements. In the peak detection algorithm above, a peak was defined as any consecutive value (photon count) that surpasses the subtracted threshold (Figure S8A_(vi)_).

In order to assess the effective concentration range of the PeCoS methodology, two-fold dilution series of Alexa488 NHS Ester were measured, and for every sample, both the ACF and the number of peaks were calculated using FCS and PeCoS (where Background = Mean ± 23*SD (water)), respectively. As expected, PeCoS did not perform well at high concentrations, where, presumably, too many particles move simultaneously through the excitation volume; at lower protein concentrations, however, (where FCS was no longer accurate) the number of peaks decreased in a nearly linear fashion (Figure S8B). To test the sensitivity of PeCoS under *in vivo* conditions, the cytoplasmic concentration of Plk4-NG was measured at the beginning of nuclear cycle 12 in embryos expressing either one (1x) or two (2x) copies of *Plk4-NG* in the *Plk4* mutant background. PeCoS analysis indicated a ~90% increase in the number of peaks (per minute) in the 2x embryos compared to the 1x embryos, indicating the effectiveness of PeCoS in measuring the relative cytosolic concentration of low abundance proteins (Figure S8C).

### Quantification and Statistical Analysis

The details for quantification, statistical tests, sample numbers, definitions of centre, and the measures for dispersion and precision are described in the main text, relevant figure legends, or relevant sections of the *Materials and Methods*. Significance in statistical tests was defined by P < 0.05. To determine whether the data values were normally distributed, a D’Agostino–Pearson omnibus normality test was applied. Prism 7 was used for all the modelling and statistical analyses.

## Supplementary Figure Legends

**Figure S1.**
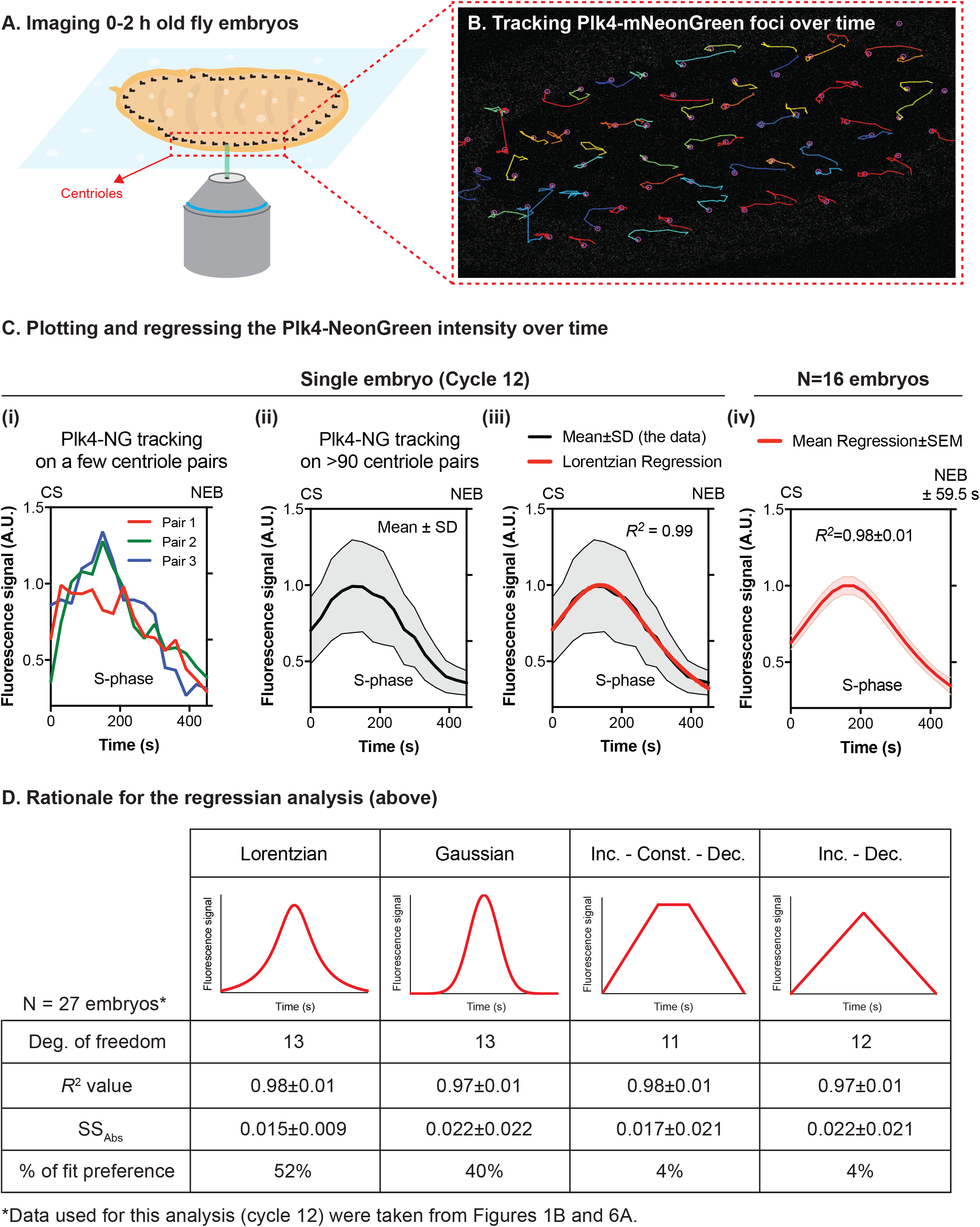
Summary of the protocol for image acquisition, processing and analysis of the Plk4-NG oscillations. (Related to Figure 1) **(A)** Diagram illustrates the centrioles in ~2 h old embryo expressing Plk4-NG being imaged on a spinning-disk confocal system. **(B)** Micrograph shows a typical image of the tracks of the Plk4-NG centrioles in S-phase of cycle 12, tracked using the ImageJ plugin, TrackMate. **(C)** Graphs show the Plk4-NG oscillation during cycle 12 in a single embryo quantified from the tracks of either several individual centriole pairs (i), or the Mean±SD oscillation calculated from the tracks of >90 centriole pairs (ii). The data for each embryo was then regressed using a Lorentzian equation (*red* line, iii)—see (D) for an explanation of the rationale for choosing this function. This process was repeated for multiple embryos to calculate a Mean±SEM regression for nuclear cycle 12 (iv). *R^2^* values indicate the goodness-of-fit (Mean±SD) of the regression. CS=time of centrosome separation (set to 0); NEB=time of nuclear envelope breakdown. **(D)** Table shows the various models that were tested to fit the Plk4-NG oscillation data. *R^2^* and *SS_Abs_* (absolute sum of squares) values indicate the goodness of fit. The Lorentzian function was the best fit for the majority of embryos, so it was used for all further analyses. Further details of these models are provided in *Materials* and *Methods*.

**Figure S2.**
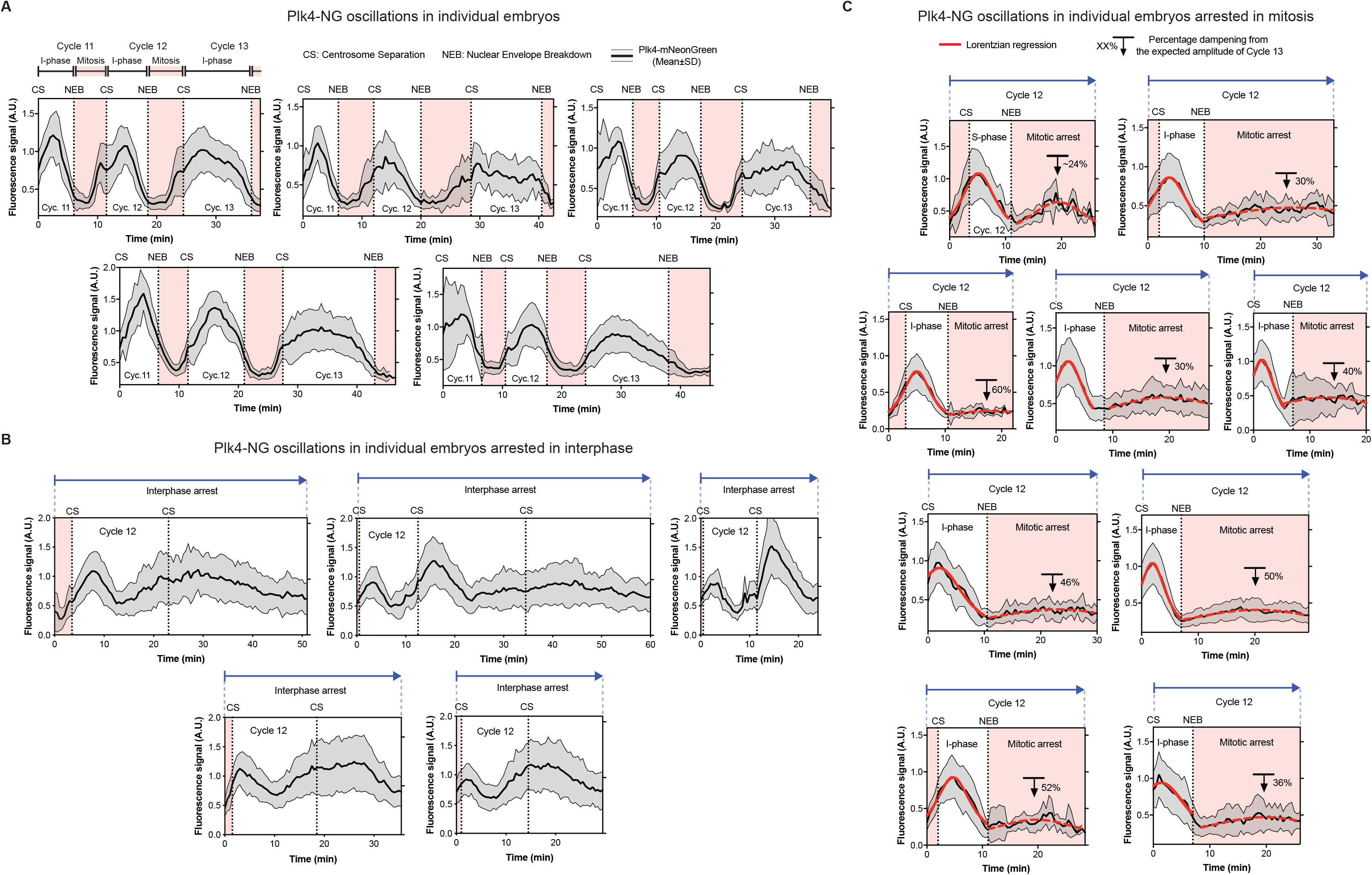
Plk4-NG oscillations in individual embryos. (Related to Figures 1, 4 and S5) **(A)** Graphs show the Mean±SD centriolar fluorescence intensity of Plk4-NG (two copies of a transgene expressed from its own promoter in a *Plk4* null mutant background) during nuclear cycles 11-13 in 5 different embryos imaged on a spinning-disk confocal system. n=26 centrioles (mean) tracked starting from cycle 11 per embryo. See *Materials* and *Methods* for full details of image acquisition and data analysis. **(B)** Same as in (A), but showing the Plk4-NG oscillation in 6 embryos arrested in interphase by the injection of dsRNAs that knock-down the levels of Cyclin A, B and B3. See Figure 4A for further details on sample numbers and experimental protocol. **(C)** Same as in (A), but showing the Plk4-NG oscillation in 9 embryos arrested in mitosis by the injection of colchicine. The first and second oscillations in these embryos were fitted independently by mathematical regression (*red* lines and *red dotted* lines, respectively). See Figure S5A for further details on the sample numbers and experimental protocol.

**Figure S3.**
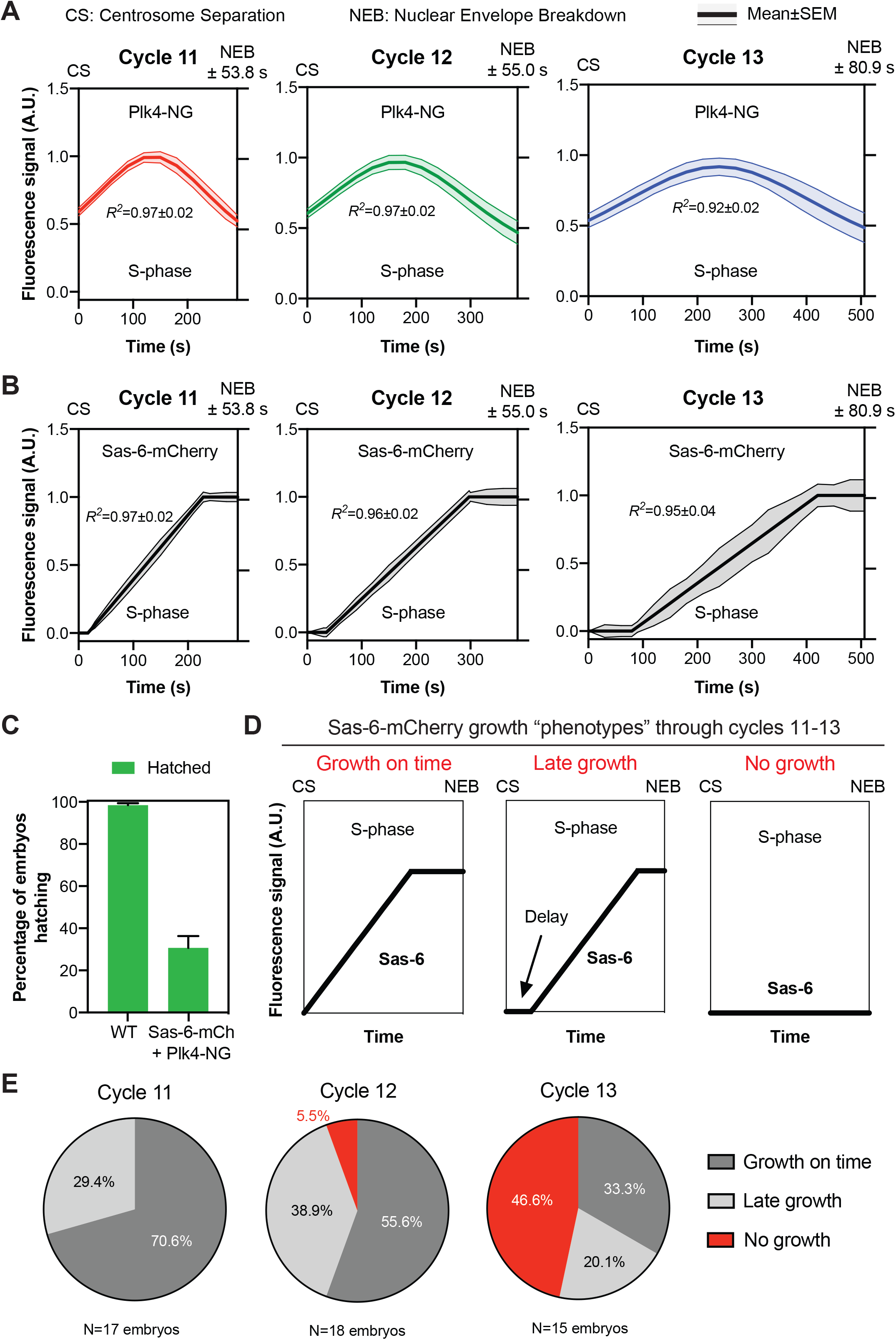
Simultaneously measuring centriole growth and the Plk4 oscillation in the same embryos. (Related to Figure 2) **(A and B)** Graphs show the same data presented in Figure 2A, but with the SEM included (as these error bars were omitted from Figure 2A for ease of presentation). CS=centrosome separation and NEB=nuclear envelope breakdown. *R^2^* values indicate the goodness of fit. **(C)** Graph quantifies the embryo hatching frequency in embryos laid by either wild-type (Oregon-R) females or females simultaneously expressing Sas-6-mCherry and Plk4-NG in a *Plk4* mutant background (all mated with WT males). At least 4 technical repeats were carried out over several days, and a total of at least 400 embryos were analysed. **(D)** Cartoon graphs (i.e. imaginary data) illustrate the three different centriole growth phenotypes we observed in the *Plk4* mutant embryos that simultaneously express 2 copies of Plk4-NG and one copy of Sas-6-mCherry. In our previous analysis of centriole growth kinetics (Aydogan et al., 2018) almost all embryos started to incorporate Sas-6-GFP at the very start of S-phase (“Growth on time”, left graph). In the embryos analysed here (with a more complicated genotype, and expressing Sas-6-mCherry rather than Sas-6-GFP), some of the embryos exhibited a clear delay in initiating the incorporation of Sas-6-mCherry (“Late growth”, middle graph), while others did not appear to incorporate significant amounts of Sas-6-mCherry at all (“No growth”, right graph); as described in the main text, many of these embryos failed to hatch (presumably due to the failure to grow their centrioles). **(E)** Pie charts quantify the percentage of embryos exhibiting each centriole growth phenotype at each nuclear cycle. Note that embryos exhibiting the “No growth” phenotype were excluded from the analysis shown in (A) and (B) and in Figure 2A, although the amplitude of the Plk4 oscillations in these embryos was analysed separately (Figure 2C): we observed 8 embryos in total that exhibited the “No growth” phenotype (1 in cycle 12, and 7 in cycle 13). Centriolar Plk4-NG levels continued to oscillate in these embryos, and the scatter graph shown in Figure 2C plots the peak amplitude of the Plk4-NG oscillations in these 8 embryos overlaid on the average “threshold” level of Plk4-NG at which centrioles started to grow in the population of embryos that did exhibit Sas-6-mCherry incorporation. This threshold was very similar at cycle 12 and 13, so the threshold shown in Figure 2C is taken from cycle 13 embryos (as 7 of the 8 embryos shown here were at cycle 13). The Plk4-NG oscillation in all but one of the 8 embryos failed to reach the average “threshold” level that would normally initiate centriole growth in these embryos.

**Figure S4.**
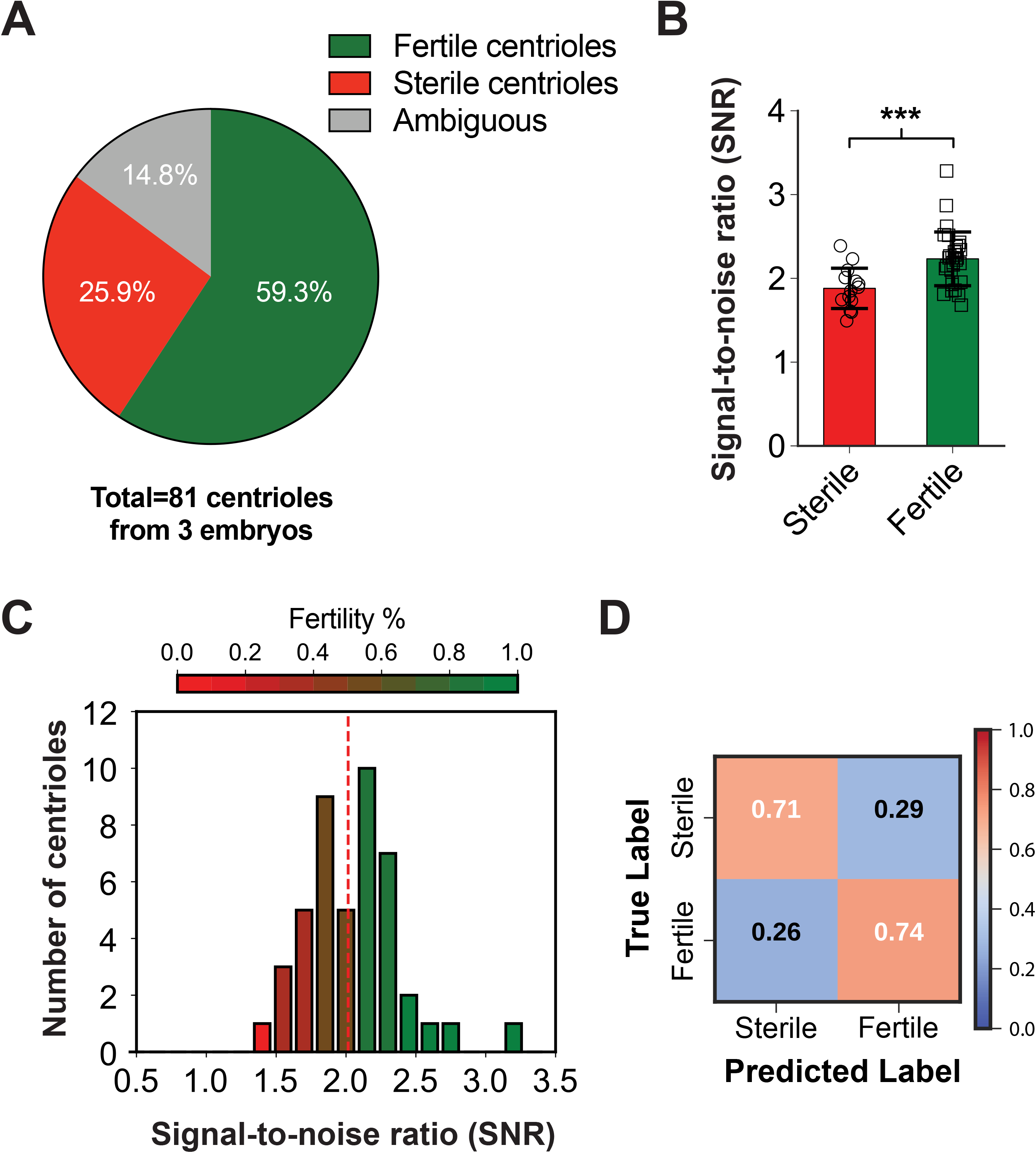
The average centriolar Plk4-NG level on individual centrioles can be used to predict centriole fertility/sterility in embryos arrested in interphase by mitotic cyclin depletion. (Related to Figures 4 and S2) **(A)** The pie chart quantifies the percentage of centrioles that continued to duplicate in embryos where cyclin A-B-B3 dsRNA was injected into embryos at nuclear cycle 2-4, and centriole behaviour assessed ~90 min later. *Ambiguous* (grey) indicates the fraction of centrioles whose duplication state could not be unambiguously determined due to their drifting out of focus during imaging. **(B)** Bar chart shows the mean signal-to-noise ratio (SNR) of Plk4-NG fluorescence signals from sterile and fertile centrioles (*red* and *green*, respectively) through the entire period of observation. Data are presented as Mean±SD. Statistical significance of SNR was tested using a *t*-test assuming equal variance (***, P<0.001). **(C)** Heatmap histogram of all SNR values from sterile and fertile centrioles. *Red dashed* line shows the unbiased threshold, determined automatically from Otsu thresholding for distinguishing sterile and fertile centrioles. Heatmap (*Red*: Sterile and *Green*: Fertile) indicates the fraction of fertile/sterile centrioles in each column. Note that, the higher the SNR, the more fertile the centrioles are. **(D)** Confusion matrix shows the classification performance of sterile versus fertile centriole Plk4-NG signals using the Otsu threshold in (C) as a proportion of the total number of signals, n=81 centrioles from 3 embryos.

**Figure S5.**
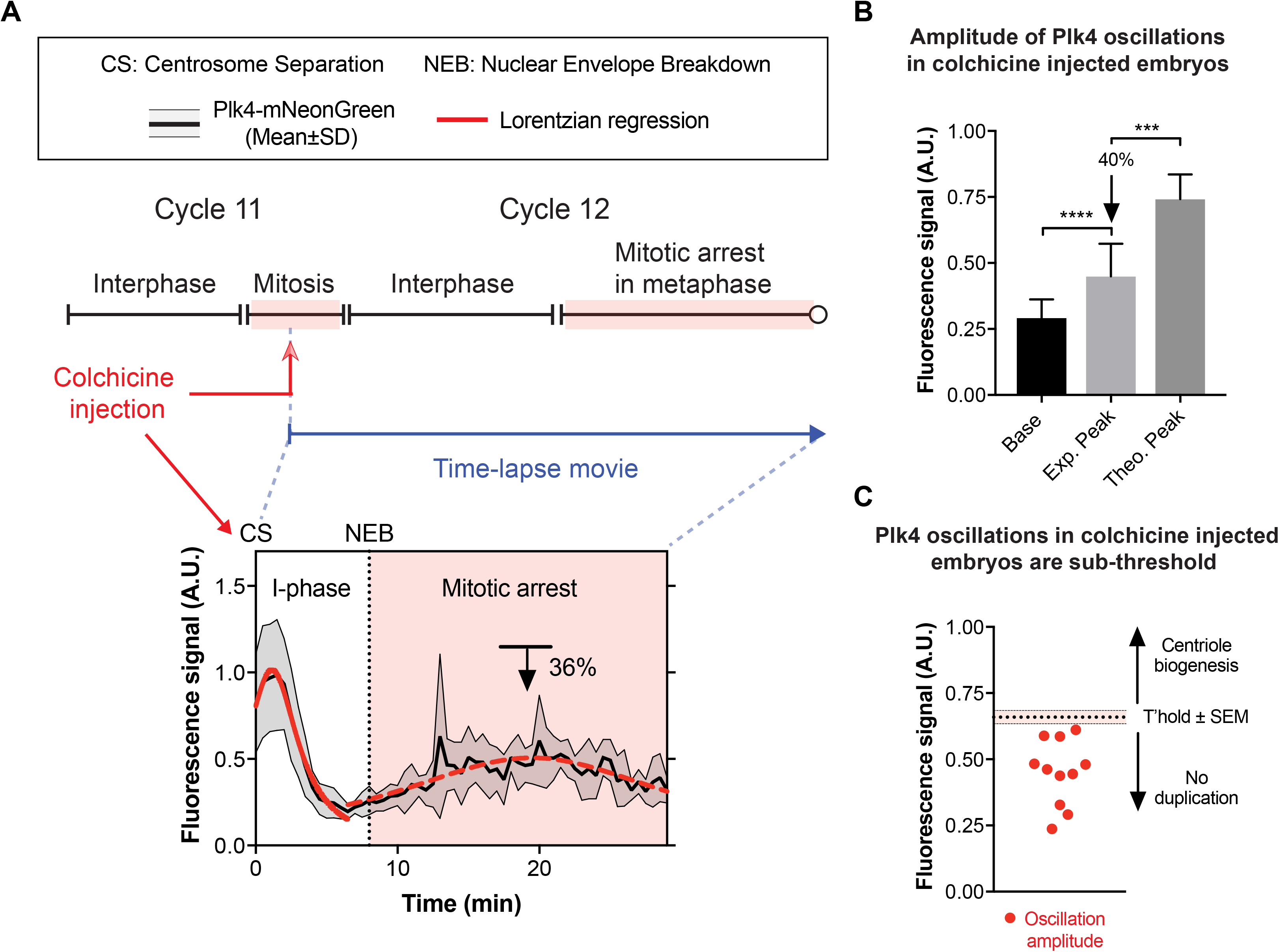
Embryos arrested in mitosis by colchicine injections exhibit Plk4 oscillations that are sub-threshold for centriole biogenesis. (Related to Figures 4 and S2) **(A)** Graph shows the Plk4-NG oscillation (*black* line, Mean±SD) in an embryo injected with colchicine at the start of S-phase of nuclear cycle 12. This embryo arrests in the next mitosis and the centrioles do not detectably duplicate again, even though a Plk4 oscillation of reduced amplitude is detectable. The first and second oscillations in these embryos were fitted by mathematical regression (*red* lines and *red dotted* lines, respectively), so that the reduction in amplitude (compared to the average amplitude in uninjected controls in cycle 13; ~36% in the embryo analysed here, *black* arrow) could be calculated. See Figure S2C for 9 additional examples. n=34 centrioles (mean) per embryo. **(B)** Bar chart illustrates that the amplitude of the second Plk4-NG oscillation observed after colchicine injection has a statistically significant difference between the base and the peak of the oscillations. On average, however, this peak is ~40% lower in amplitude than the expected mean amplitude (Theoretical Peak) of the normal Plk4-NG oscillation in cycle 13 (calculated from Figure 1B). Data are presented as Mean±SD. Statistical significance was assessed using an unpaired *t* test with Welch’s correction (for Gaussian-distributed data) or an unpaired Mann-Whitney test (***, P<0.001; ****, P<0.0001). **(C)** Scatter chart shows that the amplitude of the Plk4-NG oscillations observed during the mitotic arrest in all 11 colchicine injected embryos are below the normal threshold for centriole biogenesis in cycle 13 (threshold adapted and rescaled from the cycle 13 data shown in Figure 2).

**Figure S6.**
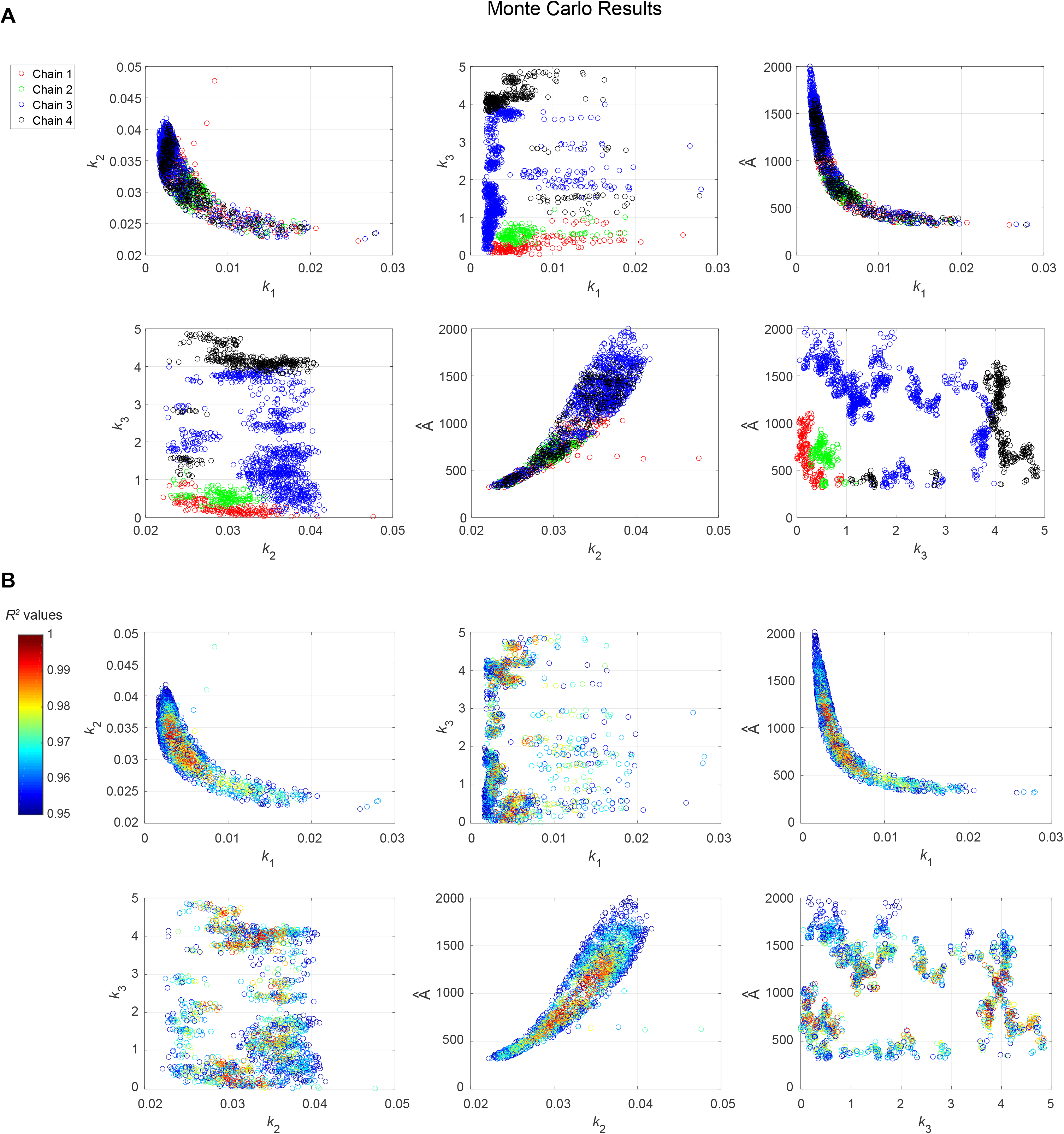
Monte Carlo simulations to characterise the parameter space of the mathematical model. (Related to Figure 5 and Table S7) **(A)** Scatter plots show the results of four Markov chain Monte Carlo simulations (shown in different colours, as indicated). The six two-dimensional projections of the four-dimensional parameter space shown here show only those points that allowed the model to fit the data well (*R^2^*>0.95). **(B)** Scatter plots show the same data points as (A), but heat-mapped to show the *R^2^* value of each point. As in (A), only the data points with *R^2^*>0.95 are shown. Points with a low *R^2^* value are shown in cool colours, while points with a high *R^2^* value are shown in warm colours (for the definition of parameters, see the mathematical modelling section in *Materials and Methods*). These results indicate that most of the parameter values that allow the model to fit the data are likely to occupy a single, continuous, and relatively small region of the parameter space. The only exception is *k_3_;* the likely reasons for this are also discussed in *Materials and Methods*.

**Figure S7.**
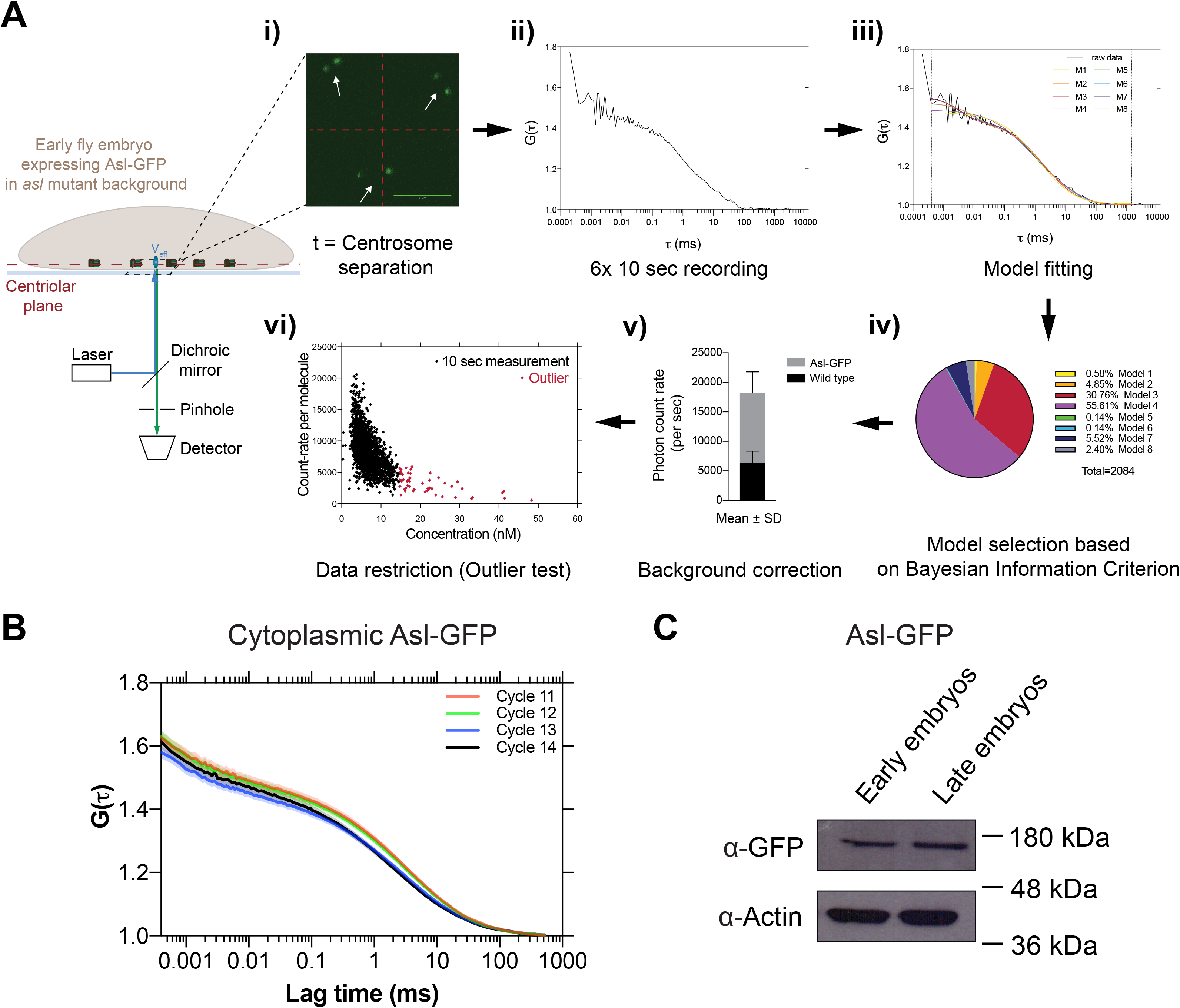
FCS analysis of cytosolic Asl levels. (Related to Figure 5) **(A)** Schematic workflow describes the acquisition and analysis of point Fluorescence Correlation Spectroscopy (FCS) measurements (see *Materials and Methods* for further details). The 488nm laser beam is positioned at the centriolar plane in embryos expressing 2 copies of Asl-GFP (under the control of its own promoter in an *asl* mutant background). **(i)** At the beginning of every cycle, when the old and new mother centrioles have just separated (*white* arrows), 6x 10sec FCS measurements were taken at a point in the cytosol maximally distant from the centrioles (centre of *red* crosshairs). **(ii)** This generated 6 autocorrelation functions (ACFs) (a typical example is shown here). **(iii)** In the FoCuS-point software, 8 different models were fitted to each ACF. **(iv)** The model that best fitted the majority of the data (#4 in this case) was chosen based on the Bayesian information criterion, and all ACFs were then fitted to this model. **(v)** The fitted ACFs were corrected for background noise which was determined by measurements in WT embryos. **(vi)** The ACFs used for further analysis were then restricted by excluding individual outlier measurements based on a ROUT-outlier test (Q = 1%) (these outlier measurements usually had a poor signal-to-noise ratio and gave concentrations that were often biologically unrealistic, and were presumably generated when a centriole or non-specific fluorescent structure passed through the analysed volume). **(B)** Graph shows the average ACFs (represented as Mean±SEM) for nuclear cycles 11-14 before background corrections. All individual ACFs were used to calculate the cytosolic concentration data shown in Figure 5E. **(C)** Western blot shows the protein levels of the Asl-GFP in either the early or late cell cycles from embryos of the same genotype used in (A) and (B). This supports the results obtained from the FCS measurements, and suggests that total Asl levels do not change significantly during the development of the syncytial embryo. Early and late embryos were separated based on their distinct morphology (judged by eye using a dissection microscope). Actin is shown as a loading control. A representative blot is shown from two technical repeats.

**Figure S8.**
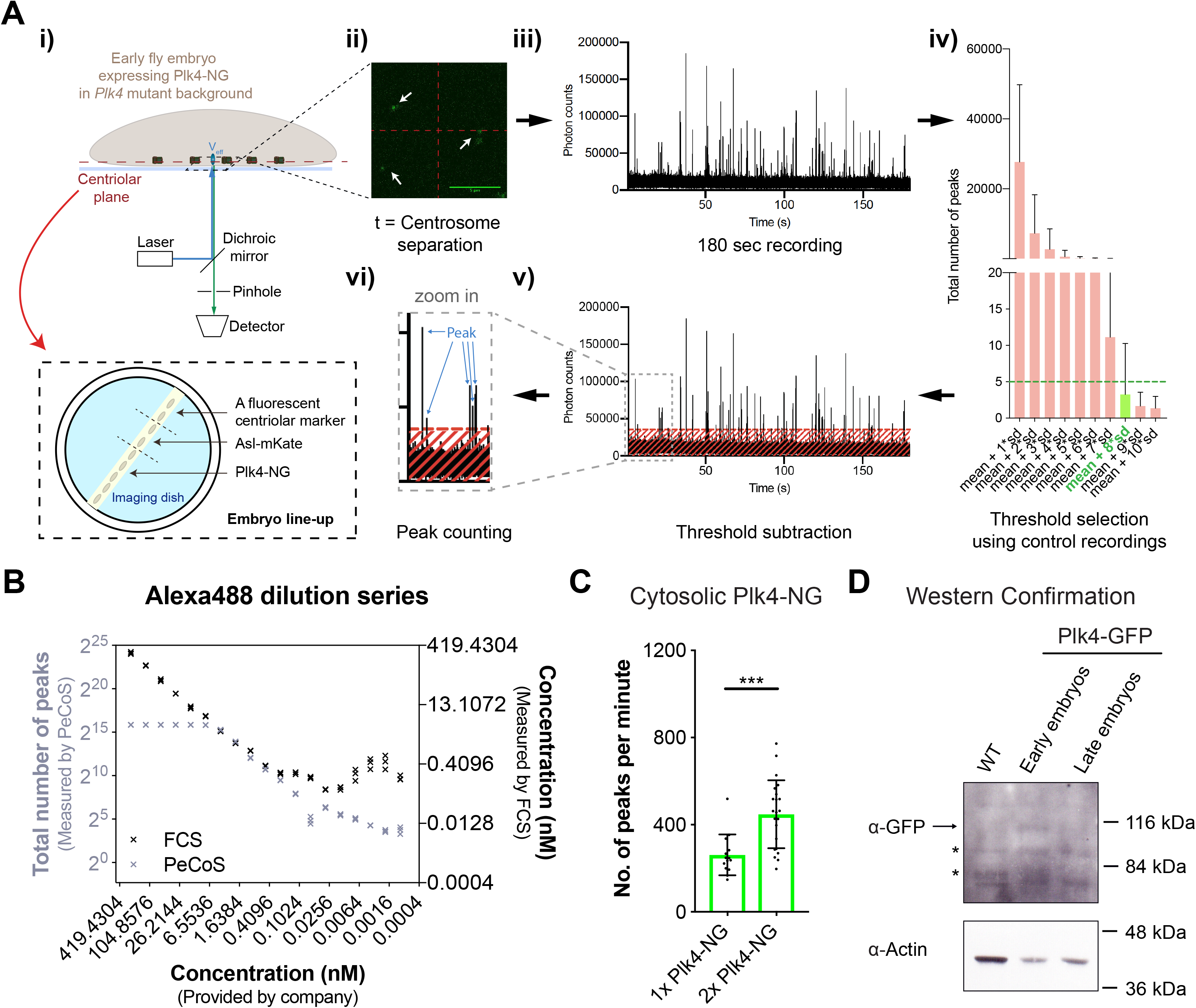
Peak Counting Spectroscopy (PeCoS) analysis of cytosolic Plk4 levels. (Related to Figure 5) **(A)** Schematic workflow describes the acquisition and analysis of *Peak Counting Spectroscopy* (PeCoS) measurements. **(i)** In addition to embryos expressing Plk4-NG under its own endogenous promoter, embryos of two other genotypes were placed on the same imaging dish. One expressing a fluorescent centriole marker to allow the spherical aberration correction caused by coverslip thickness variation, the other expressing Asl-mKate to determine the autofluorescence background threshold for the Plk4-NG expressing embryos—Asl-mKate allows one to determine the centriole plane (*white* arrows) for making these background measurements, while the red fluorophore does not interfere with the PeCoS measurements. **(ii)** Identical to FCS, a 488nm laser beam is positioned near the cortex of embryos, and the measurements are taken at a single point in the cytosol (*red* crosshairs) at the beginning of S-phase, but for 1x 180 sec **(iii)**. Afterwards, **(iv)** the threshold is calculated from control embryos, so that the background contributes less than 5 peaks on average during each recording. Following the background subtraction, **(v and vi)** the number of peaks is quantified. **(B)** To compare the effective linear concentration range of FCS and PeCoS we assessed a twofold dilution series of the Alexa488 dye. At high dye concentrations, FCS (*black* symbols) exhibits a near-linear response, while PeCoS (*grey* symbols) is saturated—presumably because there are too many fluorophores in the effective volume (*V_eff_*) for them to be measured as individual peaks. At intermediate dye concentrations, both methods exhibit a near linear response. At low concentrations (~<0.2nM), however, FCS becomes unreliable while PeCoS continues to have a near-linear response. **(C)** The bar chart shows the *in vivo* validation of PeCoS. A significant difference in the number of peaks per minute was observed between embryos expressing either 1x or 2x copies of Plk4-NG (under the control of its endogenous promoter), which were measured at the beginning of S-phase in nuclear cycle 12. Every data point represents an individual 180 sec recording. Statistical significance was assessed using an ordinary unpaired t-test (for Gaussian-distributed data) or a Mann-Whitney test (***, P<0.001). **(D)** Western blot analysis of Plk4-GFP (arrow) levels in early and late embryos supports the conclusion from the PeCoS analysis (Figure 5F), that cytosolic Plk4 levels are lower in late embryos than in early embryos. Prominent non-specific bands are indicated (*). A representative blot was shown from two technical repeats. Data are presented Mean±SD.

**Figure S9.**
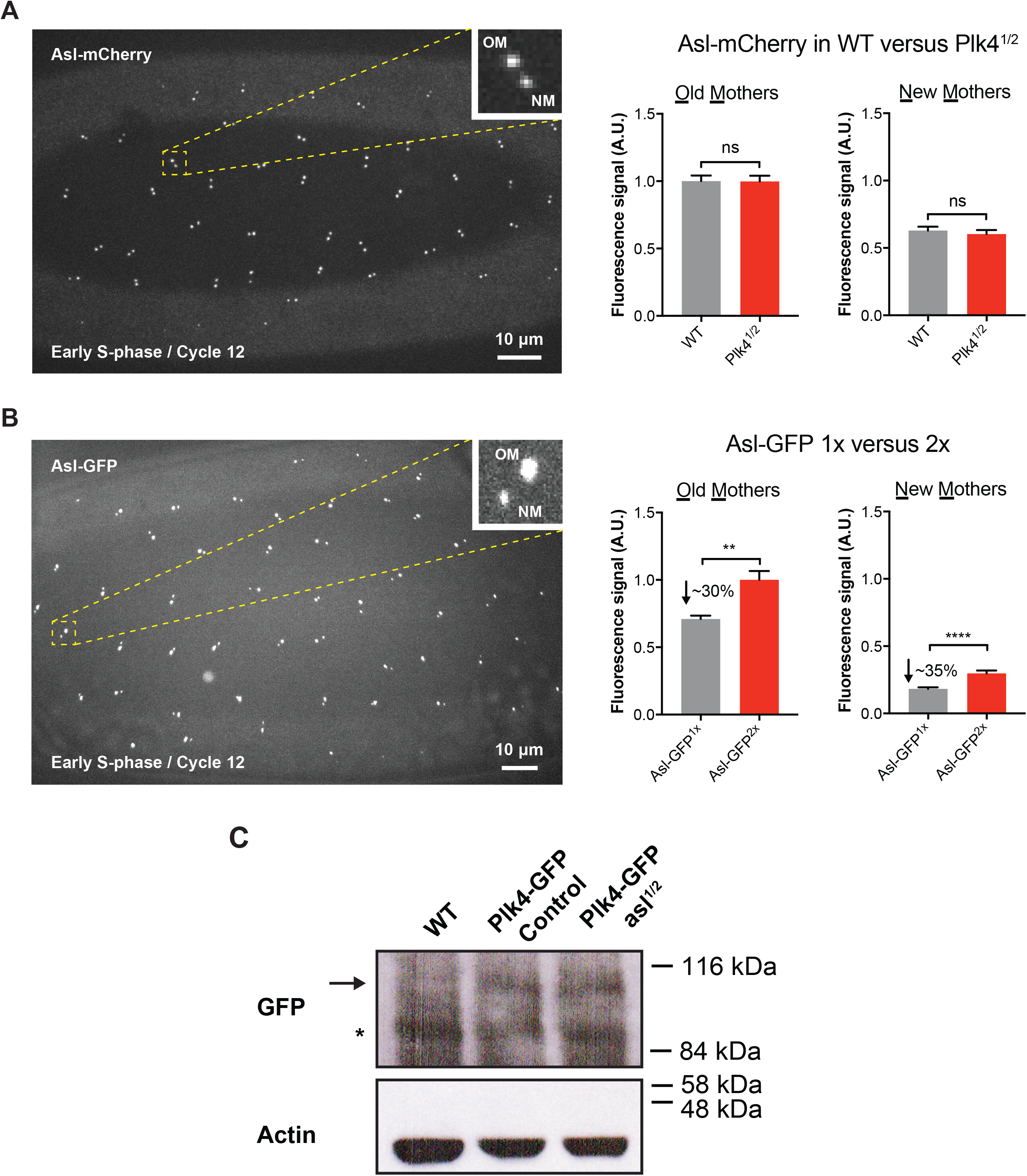
Quantification of centriolar Asl and cytoplasmic Plk4 levels when the genetic dose of *asl* or *Plk4* is halved. (Related to Figure 6) **(A)** Micrograph shows an image of Asl-mCherry at centrioles in an embryo in early S-phase (just after centrosome separation). More Asl is present on the older mother centriole (OM) than the new mother centriole (NM), as shown previously (Novak et al., 2014). Bar charts quantify the centriolar Asl-mCherry levels at OM and NM centrioles in early S-phase in either WT embryos (WT) or in embryos where the genetic dose of *Plk4* has been halved (*Plk4^1/2^*). N=17 embryos for each condition; n=67 and 58 centrioles (mean) per embryo in WT or *Plk4^1/2^* groups, respectively. Centriolar Asl levels do not change significantly when the genetic dosage of *Plk4* is halved, in agreement with the prediction of our model (see Table S1-S3). **(B)** Same schema as (A), but showing the localisation of Asl-GFP, and quantifying the centriolar levels of Asl-GFP in *asl* mutant embryos expressing either 1 (Asl-GFP^1x^) or 2 (Asl-GFP^2x^) copies of Asl-GFP. N=10 embryos for each condition; n=59 and 54 centrioles (mean) per embryo in Asl-GFP^1x^ or Asl-GFP^2x^ groups, respectively. This analysis reveals that centriolar Asl-GFP levels drop by ~30-35% when the genetic dosage of *Asl-GFP* is halved, in good agreement with the prediction of our model (see Table S1-S3). Data are represented as Mean±SEM. Statistical significance was assessed using an unpaired *t* test with Welch’s correction (for Gaussian-distributed data) or an unpaired Mann-Whitney test (**, P<0.01; ****, P<0.0001; ns, not significant). **(C)** Western blot compares the protein levels of Plk4-GFP (*arrow*) (expressed under the control of its own promoter in a *Plk4* mutant background) in otherwise WT embryos or in embryos in which the genetic dosage of *asl* has been halved. This analysis reveals that Plk4-GFP levels in the embryo do not change dramatically when the genetic dosage of *asl* is halved, in agreement with the prediction of our model. WT embryos (Lane 1) are shown as a negative control to demonstrate that the Plk4-GFP band is only detected in embryos expressing Plk4-GFP. Prominent non-specific bands are indicated (*). Actin is shown as a loading control. A representative blot is shown from two technical repeats.

## Supplementary Tables

**Table S1:** The best-fit *k*_1_, *k*_2_, *k*_3_, and *Â* parameter values^*^ for cycles 11-13. (Related to Figure 5C)

| Parameters | Cycle 11 | Cycle 12 | Cycle 13 |
| --- | --- | --- | --- |
| $k_1$ | 0.004762 | 0.003644 | 0.001272 |
| $k_2$ | 0.03756 | 0.03443 | 0.02164 |
| $k_3$ | 0.06906 | 0.06906 | 0.06906 |
| $\hat{A}$ | 1012 | 940 | 1030 |
| $\hat{A}_0$ | 701 | 637 | 557 |
| $A_{tot}$ | 1713 | 1577 | 1587 |
| $R^2$ | 0.998 | 0.9996 | 0.998 |

\*For the definition of parameters, see *Materials and Methods* under Mathematical modeling and Monte Carlo simulations.

**Table S2:** The best-fit *k*_1_, *k*_2_, *k*_3_, and *Â* parameter values^*^ for the Plk4^1/2^ experiment. (Related to Figure 6A)

| Parameters | Control | Plk4<sup>1/2</sup> |
| --- | --- | --- |
| $k_1$ | 0.003862 | 0.001513 |
| $k_2$ | 0.03699 | 0.02767 |
| $k_3$ | 0.06906 | 0.06906 |
| $\hat{A}$ | 4884 | 5607 |
| $\hat{A}_0$ | 3152 | 2390 |
| $A_{tot}$ | 8036 | 7997 |
| $R^2$ | 0.9992 | 0.996 |

\*For the definition of parameters, see *Materials and Methods* under Mathematical modeling and Monte Carlo simulations.

**Table S3:** The best-fit *k*_1_, *k*_2_, *k*_3_, and *Â* parameter values* for the *asl^1/2^* experiment. (Related to Figure 6B)

| Parameters | Control | asl<sup>1/2</sup> |
| --- | --- | --- |
| $k_1$ | 0.003437 | 0.002724 |
| $k_2$ | 0.03246 | 0.03324 |
| $k_3$ | 0.06906 | 0.06906 |
| $\hat{A}$ | 1938 | 1408 |
| $\hat{A}_0$ | 1298 | 950 |
| $A_{tot}$ | 3236 | 2358 |
| $R^2$ | 0.9996 | 0.9991 |

\*For the definition of parameters, see *Materials and Methods* under Mathematical modeling and Monte Carlo simulations.

**Table S4:**
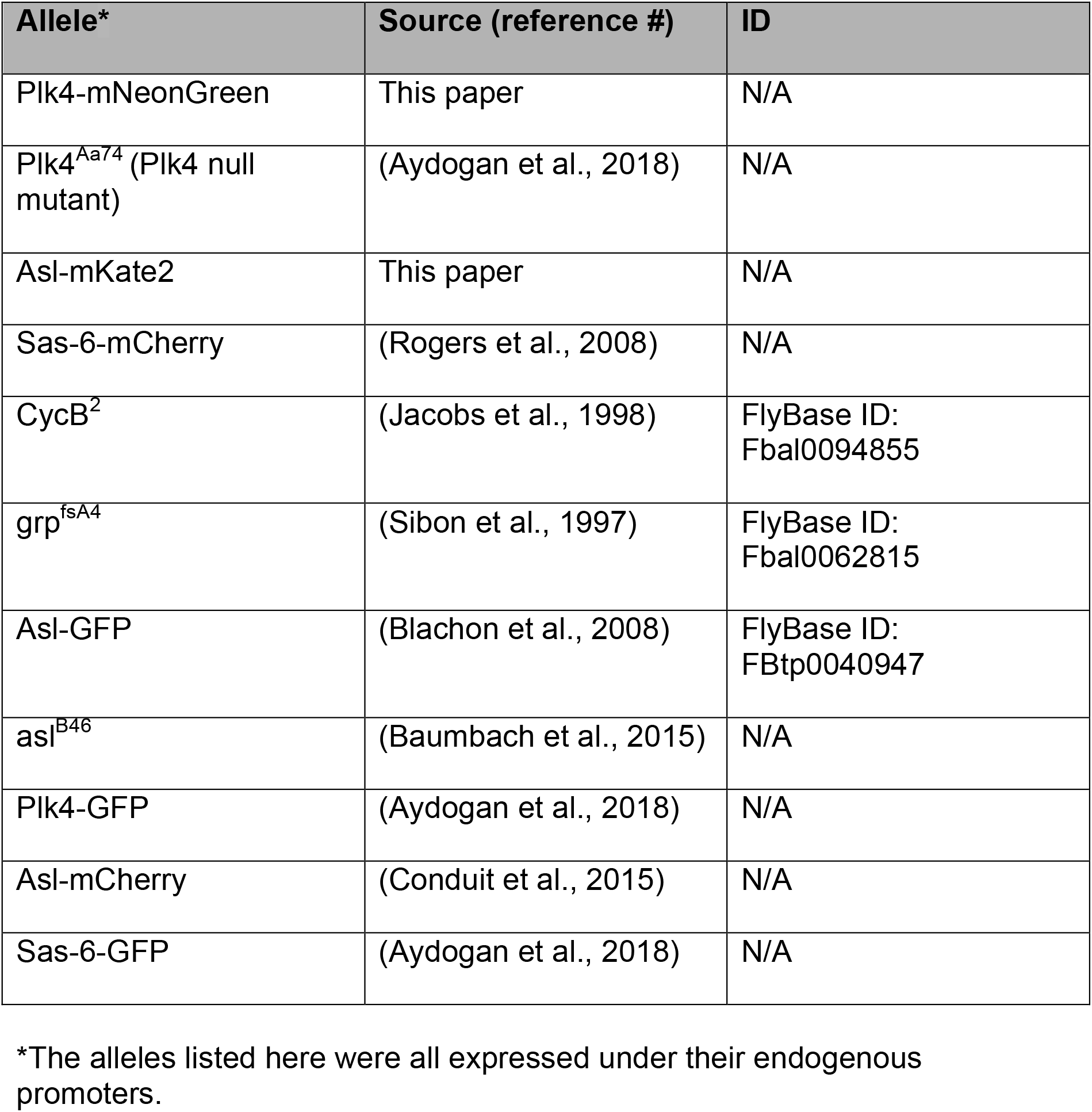
*D. melanogaster* allelles used in this study.

**Table S5:**
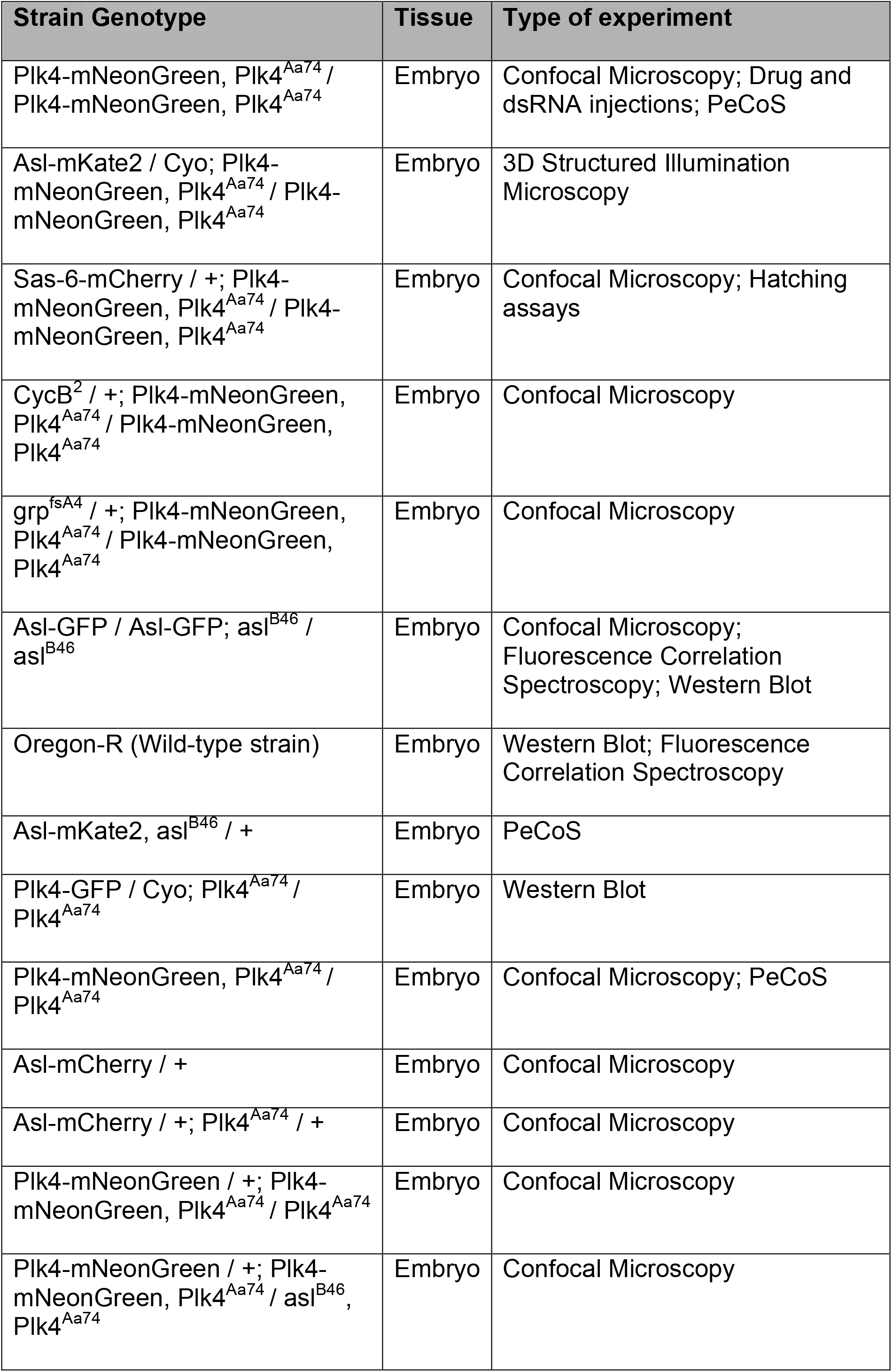
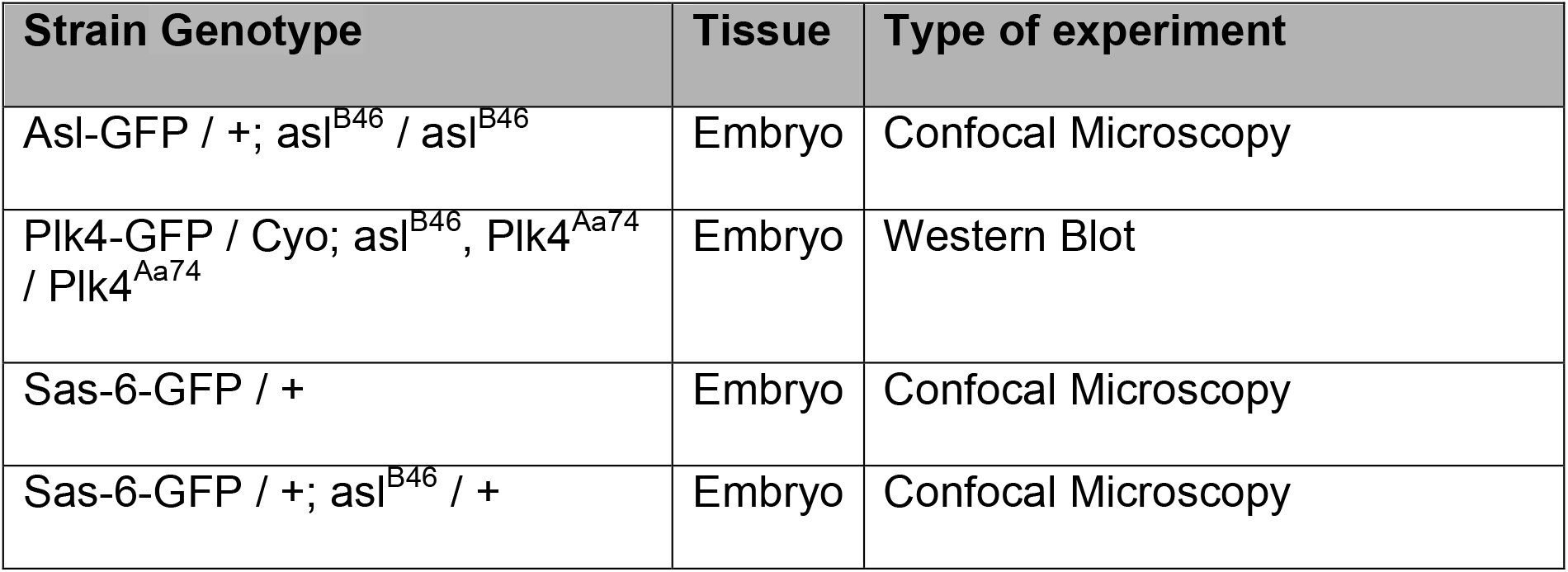
*D. melanogaster* strains generated and/or used in this study.

**Table S6:**
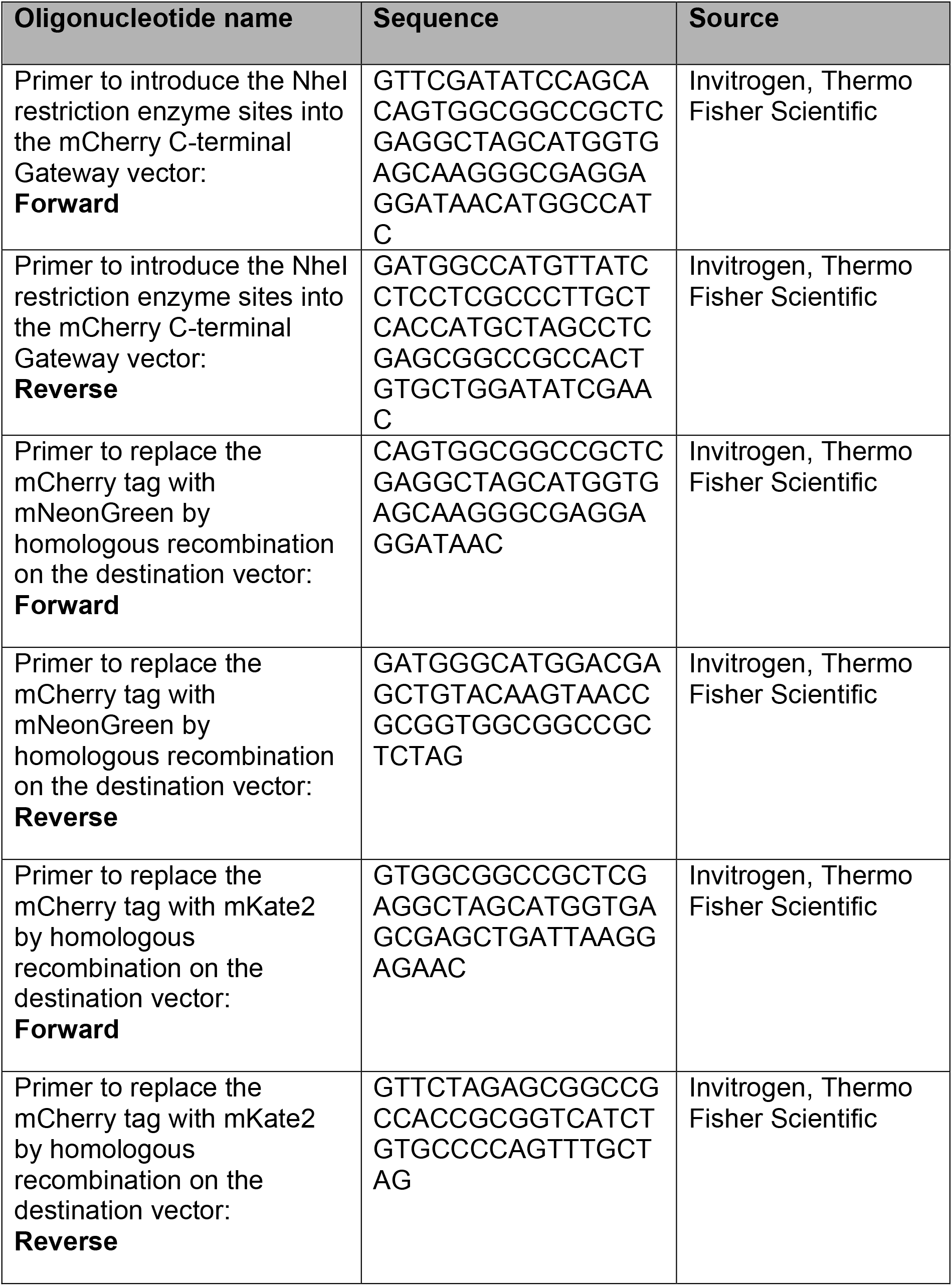
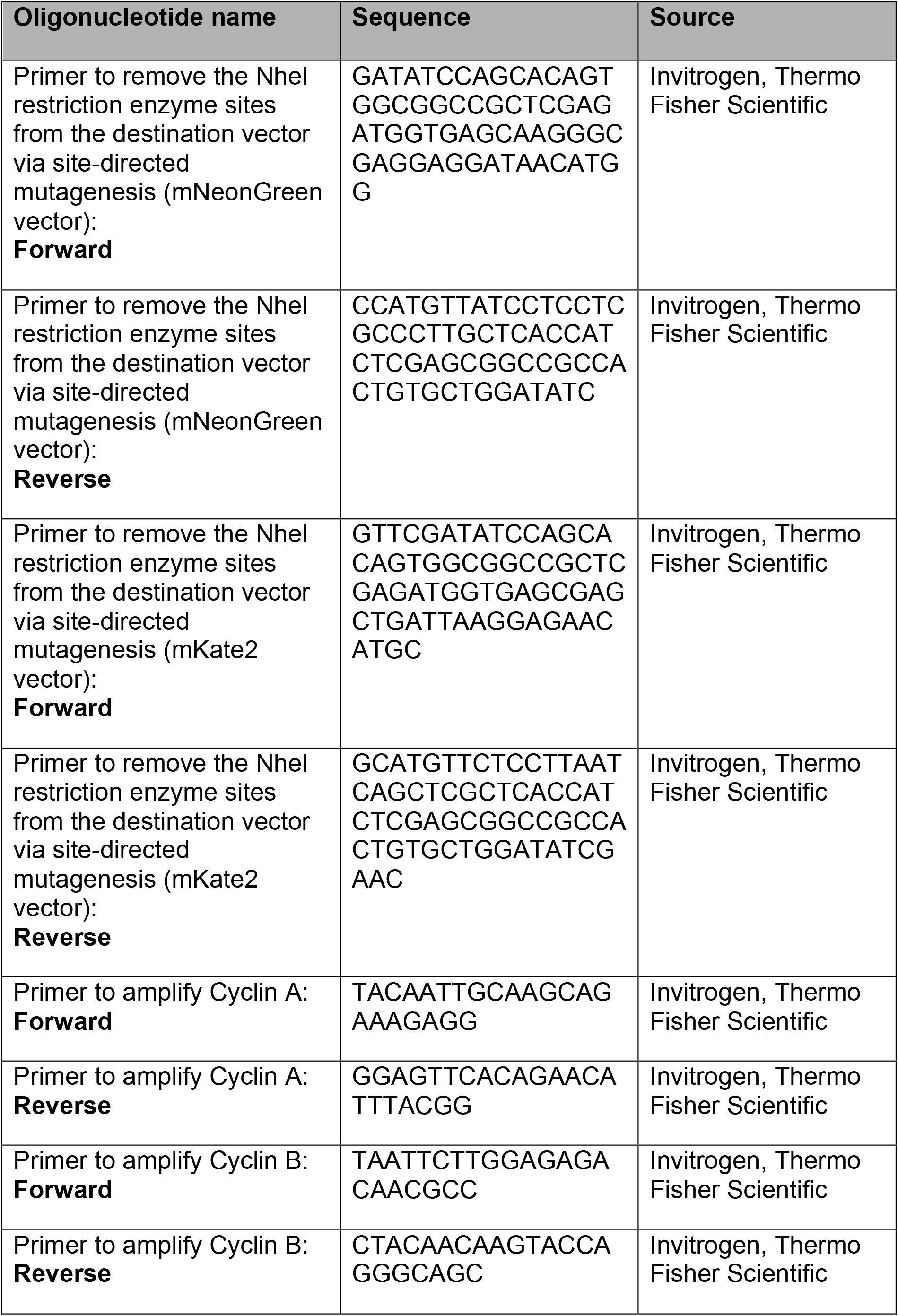
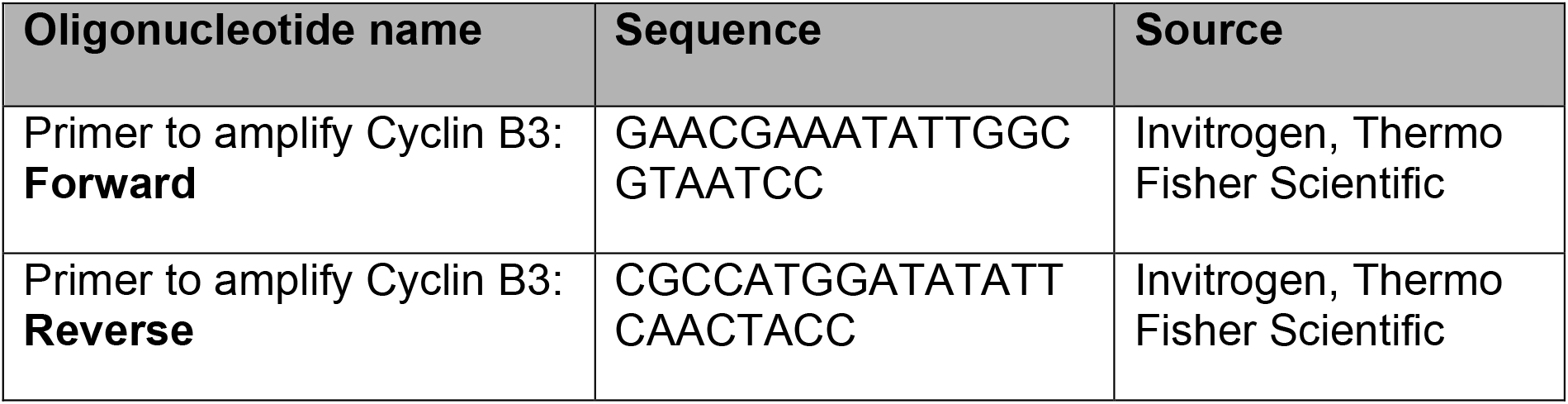
Oligonucleotides used in this study.

**Table S7:** Starting parameter values and proposal acceptance rates for each of four Markov chains used in Metropolis-Hastings Markov chain Monte Carlo. (Related to Figure S6)

| Parameters | Chain 1 | Chain 2 | Chain 3 | Chain 4 |
| --- | --- | --- | --- | --- |
| $k_1$ | 0.0035 | 0.01 | 0.1 | 0.0035 |
| $k_2$ | 0.035 | 0.09 | 0.2 | 0.035 |
| $k_3$ | 0.05 | 0.5 | 0.1 | 4 |
| $\hat{A}$ | 1000 | 500 | 1500 | 1500 |
| Acceptance | 36% | 36% | 29% | 37% |

## Supplementary Movie Captions

**Video S1. Monitoring Plk4-NG oscillations in a *Drosophila* embryo.**

Time-lapse video of an embryo expressing two copies of Plk4-NG (expressed transgenically from the endogenous *Plk4* promoter) in a *Plk4* mutant background, observed on a spinning-disk confocal microscope through nuclear cycles 11-13. The movie is a maximum-intensity projection that has been photo-bleach corrected, but not background subtracted for visual clarity. Time (min:sec) is shown at the top left, and stage of the cell cycle is indicated at the bottom left.

**Video S2. Monitoring Sas-6-mCherry incorporation and Plk4-NG oscillations simultaneously in the same embryo.**

Time-lapse movie of an embryo expressing one copy of Sas-6-mCherry (expressed transgenically from the endogenous *Sas-6* promotor) and two copies of Plk4-NG (expressed transgenically from the endogenous *Plk4* promoter) in a *Plk4* mutant background, observed on a spinning-disk confocal microscope during S-phase of nuclear cycle 12. The movie is a maximum-intensity projection that has been photo-bleach corrected, but not background subtracted for visual clarity. Time (Min:Sec) is shown at the top left, and stage of the cell cycle is indicated at the bottom left.

**Video S3. Centrioles can continue to duplicate in embryos arrested for a short period in interphase by mitotic cyclin depletion.**

Time-lapse movie of an embryo expressing two copies of Plk4-NG (expressed transgenically from the endogenous *Plk4* promoter) in a *Plk4* mutant background, observed on a spinning-disk confocal microscope through nuclear cycle 12. The embryo was injected with cyclin A-B-B3 dsRNA in ~cycle 8, approximately 30-40 min prior to the start of the movie. The movie on the left is the maximum intensity projection of the slices where centrioles are in focus. The movie on the right is the maximum intensity projection of the slices where nuclei are in focus. Note how the centrioles undergo at least two rounds of duplication, the second of which is more asynchronous and occurs without nuclear envelope breakdown (indicating that the nuclei are arrested in an interphase-like state). These videos have been photo-bleach corrected, but not background subtracted for visual clarity. Time (Min:Sec) is shown at the top left.

**Video S4. Centrioles duplicate stochastically in embryos arrested for a long period in interphase by mitotic cyclin depletion.**

Time-lapse movie of an embryo expressing two copies of Plk4-NG (expressed transgenically from the endogenous *Plk4* promoter) in a *Plk4* mutant background, observed on a spinning-disk confocal microscope. The embryo was injected with cyclin A-B-B3 dsRNA in ~cycle 2-4, approximately 90 min prior to the start of the movie. The movie on the left is the maximum intensity projection of the slices where centrioles are in focus. The movie on the right is the maximum intensity projection of the slices where nuclei are in focus. Note that a small number of large nuclei are present throughout the time-course of the movie (indicating that they are arrested in an interphase-like state), but some centrioles duplicate one or more times in an apparently stochastic manner. The Plk4-NG oscillations on individual centrioles are less obvious than in normally cycling embryos, but an unbiased computational analysis of these movies indicates that individual centriole duplication events are correlated with individual centriolar Plk4-NG oscillations (Figures 4B–E and S4). These videos have been photo-bleach corrected, but not background subtracted for visual clarity. Time (Min:Sec) is shown at the top left.

**Video S5. Centrioles do not duplicate when arrested in mitosis by colchicine injection.**

Time-lapse movie of an embryo expressing two copies of Plk4-NG (expressed transgenically from the endogenous *Plk4* promoter) in a *Plk4* mutant background, observed on a spinning-disk confocal microscope through nuclear cycle 12. The embryo was injected with colchicine in mitosis of nuclear cycle 11, approximately 30 sec prior to the start of the movie. The movie is a maximum-intensity projection that has been photo-bleach corrected, but not background subtracted for visual clarity. Time (Min:Sec) is shown at the top left, and stage of the cell cycle is indicated at the bottom left.

## References

Alvarez Rodrigo, I., Conduit, P.T., Baumbach, J., Novak, Z.A., Aydogan, M.G., Wainman, A., and Raff, J.W. (2018). A positive feedback loop drives centrosome maturation in flies. 380907.

Aydogan, M.G., Wainman, A., Saurya, S., Steinacker, T.L., Caballe, A., Novak, Z.A., Baumbach, J., Muschalik, N., and Raff, J.W. (2018). A homeostatic clock sets daughter centriole size in flies. J Cell Biol 217, 1233–1248.

Ball, G., Demmerle, J., Kaufmann, R., Davis, I., Dobbie, I.M., and Schermelleh, L. (2015). SIMcheck: a Toolbox for Successful Super-resolution Structured Illumination Microscopy. Sci Rep 5, 15915.

Banterle, N., and Gönczy, P. (2017). Centriole Biogenesis: From Identifying the Characters to Understanding the Plot. Annu Rev Cell Dev Biol 33, 23–49.

Basto, R., Brunk, K., Vinadogrova, T., Peel, N., Franz, A., Khodjakov, A., and Raff, J.W. (2008). Centrosome amplification can initiate tumorigenesis in flies. Cell 133, 1032–1042.

Baumbach, J., Novak, Z.A., Raff, J.W., and Wainman, A. (2015). Dissecting the function and assembly of acentriolar microtubule organizing centers in Drosophila cells in vivo. PLoS Genet 11, e1005261.

Bettencourt-Dias, M., Hildebrandt, F., Pellman, D., Woods, G., and Godinho, S.A. (2011). Centrosomes and cilia in human disease. Trends in Genetics 27, 307–315.

Blachon, S., Gopalakrishnan, J., Omori, Y., Polyanovsky, A., Church, A., Nicastro, D., Malicki, J., and Avidor-Reiss, T. (2008). Drosophila asterless and vertebrate Cep152 Are orthologs essential for centriole duplication. Genetics 180, 2081–2094.

Boese, C.J., Nye, J., Buster, D.W., McLamarrah, T.A., Byrnes, A.E., Slep, K.C., Rusan, N.M., and Rogers, G.C. (2018). Asterless is a Polo-like kinase 4 substrate that both activates and inhibits kinase activity depending on its phosphorylation state. Mol Biol Cell 29, 2874–2886.

Cizmecioglu, O., Arnold, M., Bahtz, R., Settele, F., Ehret, L., Haselmann-Weiβ, U., Antony, C., and Hoffmann, I. (2010). Cep152 acts as a scaffold for recruitment of Plk4 and CPAP to the centrosome. J Cell Biol 191, 731–739.

Claude, A. (1943). The constitution of protoplasm. Science 97, 451–456.

Conduit, P.T., Wainman, A., Novak, Z.A., Weil, T.T., and Raff, J.W. (2015). Re-examining the role of Drosophila Sas-4 in centrosome assembly using two-colour-3D-SIM FRAP. Elife 4, 1032.

Cunha-Ferreira, I., Bento, I., Pimenta-Marques, A., Jana, S.C., Lince-Faria, M., Duarte, P., Borrego-Pinto, J., Gilberto, S., Amado, T., Brito, D., et al. (2013). Regulation of Autophosphorylation Controls PLK4 Self-Destruction and Centriole Number. Current Biology 23, 2245–2254.

Dzhindzhev, N.S., Tzolovsky, G., Lipinszki, Z., Schneider, S., Lattao, R., Fu, J., Debski, J., Dadlez, M., and Glover, D.M. (2014). Plk4 phosphorylates Ana2 to trigger Sas6 recruitment and procentriole formation. Curr. Biol 24, 2526–2532.

Dzhindzhev, N.S., Yu, Q.D., Weiskopf, K., Tzolovsky, G., Cunha-Ferreira, I., Riparbelli, M., Rodrigues-Martins, A., Bettencourt-Dias, M., Callaini, G., and Glover, D.M. (2010). Asterless is a scaffold for the onset of centriole assembly. Nature 467, 714–718.

Eilers, P.H., and Boelens, H.F. (2005). Baseline Correction with Asymmetric Least Squares Smoothing. 1–24.

Ferrell, J.E. (2016). Perfect and Near-Perfect Adaptation in Cell Signaling. Cell Syst 2, 62–67.

Friedman, S.L. (2000). Molecular regulation of hepatic fibrosis, an integrated cellular response to tissue injury. J Biol Chem 275, 2247–2250.

Firat-Karalar, E.N., and Stearns, T. (2014). The centriole duplication cycle. Curr. Biol 369, 20130460–20130460.

Goehring, N.W., and Hyman, A.A. (2012). Organelle Growth Control through Limiting Pools of Cytoplasmic Components. Current Biology 22, R330–R339.

Guichard, P., Hachet, V., Majubu, N., Neves, A., Demurtas, D., Olieric, N., Flückiger, I., Yamada, A., Kihara, K., Nishida, Y., et al. (2013). Native architecture of the centriole proximal region reveals features underlying its 9-fold radial symmetry. 23, 1620–1628.

Hatch, E.M., Kulukian, A., Holland, A.J., Cleveland, D.W., and Stearns, T. (2010). Cep152 interacts with Plk4 and is required for centriole duplication. J Cell Biol 191, 721–729.

Holland, A.J., Lan, W., Niessen, S., Hoover, H., and Cleveland, D.W. (2010). Polo-like kinase 4 kinase activity limits centrosome overduplication by autoregulating its own stability. J Cell Biol 188, 191–198.

Jacobs, H.W., Knoblich, J.A., and Lehner, C.F. (1998). Drosophila Cyclin B3 is required for female fertility and is dispensable for mitosis like Cyclin B. Genes Dev 12, 3741–3751.

Jones, E., Oliphant, T., org, P.P.W.W.S., (2001) SciPy: Open Source Scientific Tools for Python. [Online]. Available: ht. tp.

Khodjakov, A., Rieder, C.L., Sluder, G., Cassels, G., Sibon, O., and Wang, CL. (2002). De novo formation of centrosomes in vertebrate cells arrested during S phase. J Cell Biol 158, 1171–1181.

Kitagawa, D., Vakonakis, I., Olieric, N., Hilbert, M., Keller, D., Olieric, V., Bortfeld, M., Erat, M.C., Flückiger, I., Gönczy, P., et al. (2011). Structural basis of the 9-fold symmetry of centrioles. Cell 144, 364–375.

Klebba, J.E., Buster, D.W., Nguyen, A.L., Swatkoski, S., Gucek, M., Rusan, N.M., and Rogers, G.C. (2013). Polo-like Kinase 4 Autodestructs by Generating Its Slimb-Binding Phosphodegron. Current Biology 23, 2255–2261.

Klebba, J.E., Galletta, B.J., Nye, J., Plevock, K.M., Buster, D.W., Hollingsworth, N.A., Slep, K.C., Rusan, N.M., and Rogers, G.C. (2015). Two Polo-like kinase 4 binding domains in Asterless perform distinct roles in regulating kinase stability. J Cell Biol 208, 401–414.

Koppel, D.E. (1974). Statistical accuracy in fluorescence correlation spectroscopy. Physical Review A 10, 1938–1945.

Kratz, A.-S., Bärenz, F., Richter, K.T., and Hoffmann, I. (2015). Plk4-dependent phosphorylation of STIL is required for centriole duplication. Biology Open 4, 370–377.

Leda, M., Holland, A.J., and Goryachev, A.B. (2018). Autoamplification and Competition Drive Symmetry Breaking: Initiation of Centriole Duplication by the PLK4-STIL Network. iScience 8, 222–235.

Liu, T.-L., Upadhyayula, S., Milkie, D.E., Singh, V., Wang, K., Swinburne, I.A., Mosaliganti, K.R., Collins, Z.M., Hiscock, T.W., Shea, J., et al. (2018). Observing the cell in its native state: Imaging subcellular dynamics in multicellular organisms. Science 360, eaaq1392.

Lu, Y., and Cross, F.R. (2010). Periodic cyclin-Cdk activity entrains an autonomous Cdc14 release oscillator. Cell 141, 268–279.

Marsh, B.J., Mastronarde, D.N., Buttle, K.F., Howell, K.E., and McIntosh, J.R. (2001). Organellar relationships in the Golgi region of the pancreatic beta cell line, HIT-T15, visualized by high resolution electron tomography. Proceedings of the National Academy of Sciences 98, 2399–2406.

Marshall, W.F. (2016). Cell Geometry: How Cells Count and Measure Size. Annu Rev Biophys 45, 49–64.

Matsuo, T., Yamaguchi, S., Mitsui, S., Emi, A., Shimoda, F., and Okamura, H. (2003). Control mechanism of the circadian clock for timing of cell division in vivo. Science 302, 255–259.

McCleland, M.L., and O’Farrell, P.H. (2008). RNAi of Mitotic Cyclins in Drosophila Uncouples the Nuclear and Centrosome Cycle. Curr. Biol 18, 245–254.

Mukherji, S., and O’Shea, E.K. (2014). Mechanisms of organelle biogenesis govern stochastic fluctuations in organelle abundance. Elife 3, e02678.

Nigg, E.A., and Holland, A.J. (2018). Once and only once: mechanisms of centriole duplication and their deregulation in disease. Nat Rev Mol Cell Biol 19, 297–312.

Nigg, E.A., and Raff, J.W. (2009). Centrioles, centrosomes, and cilia in health and disease. Cell 139, 663–678.

Novak, Z.A., Conduit, P.T., Wainman, A., and Raff, J.W. (2014). Asterless licenses daughter centrioles to duplicate for the first time in Drosophila embryos. Curr. Biol. 24, 1276–1282.

Novak, Z.A., Wainman, A., Gartenmann, L., and Raff, J.W. (2016). Cdk1 Phosphorylates Drosophila Sas-4 to Recruit Polo to Daughter Centrioles and Convert Them to Centrosomes. Dev Cell 37, 545–557.

Novák, B., and Tyson, J.J. (2008). Design principles of biochemical oscillators. Nat Rev Mol Cell Biol 9, 981–991.

Ohta, M., Ashikawa, T., Nozaki, Y., Kozuka-Hata, H., Goto, H., Inagaki, M., Oyama, M., and Kitagawa, D. (2014). Direct interaction of Plk4 with STIL ensures formation of a single procentriole per parental centriole. Nat Commun 5, 5267.

Ohta, M., Watanabe, K., Ashikawa, T., Nozaki, Y., Yoshiba, S., Kimura, A., and Kitagawa, D. (2018). Bimodal Binding of STIL to Plk4 Controls Proper Centriole Copy Number. Cell Rep 23, 3160–3169.e3164.

Orlando, D.A., Lin, C.Y., Bernard, A., Wang, J.Y., Socolar, J.E.S., Iversen, E.S., Hartemink, A.J., and Haase, S.B. (2008). Global control of cell-cycle transcription by coupled CDK and network oscillators. Nature 453, 944–947.

Petrášek, Z., and Schwille, P. (2008). Precise Measurement of Diffusion Coefficients using Scanning Fluorescence Correlation Spectroscopy. Biophys J 94, 1437–1448.

Raff, J.W., and Glover, D.M. (1989). Centrosomes, and not nuclei, initiate pole cell formation in Drosophila embryos. Cell 57, 611–619.

Rahi, S.J., Pecani, K., Ondracka, A., Oikonomou, C., and Cross, F.R. (2016). The CDK-APC/C Oscillator Predominantly Entrains Periodic Cell-Cycle Transcription. Cell 165, 475–487.

Rogers, G.C., Rusan, N.M., Peifer, M., and Rogers, S.L. (2008). A multicomponent assembly pathway contributes to the formation of acentrosomal microtubule arrays in interphase Drosophila cells. Mol Biol Cell 19, 3163–3178.

Rüttinger, S., Buschmann, V., Krämer, B., Erdmann, R., Macdonald, R., and Koberling, F. (2008). Comparison and accuracy of methods to determine the confocal volume for quantitative fluorescence correlation spectroscopy. J Microsc 232, 343–352.

Schönle, A., Middendorff, Von. C., Ringemann, C., Hell, S.W., and Eggeling, C. (2014). Monitoring triplet state dynamics with fluorescence correlation spectroscopy: bias and correction. Microsc Res Tech 77, 528–536.

Schwarz, G. (1978). Estimating the Dimension of a Model. The Annals of Statistics 6, 461–464.

Shaner, N.C., Lambert, G.G., Chammas, A., Ni, Y., Cranfill, P.J., Baird, M.A., Sell, B.R., Allen, J.R., Day, R.N., Israelsson, M., et al. (2013). A bright monomeric green fluorescent protein derived from Branchiostoma lanceolatum. Nat Methods 10, 407–409.

Shcherbo, D., Murphy, C.S., Ermakova, G.V., Solovieva, E.A., Chepurnykh, T.V., Shcheglov, A.S., Verkhusha, V.V., Pletnev, V.Z., Hazelwood, K.L., Roche, P.M., et al. (2009). Far-red fluorescent tags for protein imaging in living tissues. Biochem. J. 418, 567–574.

Sibon, O.C., Stevenson, V.A., and Theurkauf, W.E. (1997). DNA-replication checkpoint control at the Drosophila midblastula transition. Nature 388, 93–97.

Somvanshi, P.R., Patel, A.K., Bhartiya, S., and Venkatesh, K.V. (2015). Implementation of integral feedback control in biological systems. Wiley Interdiscip Rev Syst Biol Med 7, 301–316.

Stern, B., and Nurse, P. (1996). A quantitative model for the cdc2 control of S phase and mitosis in fission yeast. Trends Genet. 12, 345–350.

Swaffer, M.P., Jones, A.W., Flynn, H.R., Snijders, A.P., and Nurse, P. (2016). CDK Substrate Phosphorylation and Ordering the Cell Cycle. Cell 167, 1750–1761.e16.

Swaffer, M.P., Jones, A.W., Flynn, H.R., Snijders, A.P., and Nurse, P. (2018). Quantitative Phosphoproteomics Reveals the Signaling Dynamics of Cell-Cycle Kinases in the Fission Yeast Schizosaccharomyces pombe. Cell Rep 24, 503–514.

Tinevez, J.-Y., Perry, N., Schindelin, J., Hoopes, G.M., Reynolds, G.D., Laplantine, E., Bednarek, S.Y., Shorte, S.L., and Eliceiri, K.W. (2016). TrackMate: An open and extensible platform for single-particle tracking. Methods 115, 80–90.

van Breugel, M., Hirono, M., Andreeva, A., Yanagisawa, H.-A., Yamaguchi, S., Nakazawa, Y., Morgner, N., Petrovich, M., Ebong, I.-O., Robinson, C.V., et al. (2011). Structures of SAS-6 suggest its organization in centrioles. Science 331, 1196–1199.

van Breugel, M., Wilcken, R., McLaughlin, S.H., Rutherford, T.J., and Johnson, C.M. (2014). Structure of the SAS-6 cartwheel hub from Leishmania major. Elife 3, e01812.

Vidwans, S.J., Wong, M.L., and O’Farrell, P.H. (2003). Anomalous centriole configurations are detected in Drosophila wing disc cells upon Cdk1 inactivation. J Cell Sci 116, 137–143.

Waithe, D., Clausen, M.P., Sezgin, E., and Eggeling, C. (2016). FoCuS-point: software for STED fluorescence correlation and time-gated single photon counting. Bioinformatics 32, 958–960.

Wang, Y., Song, L., Liu, M., Ge, R., Zhou, Q., Liu, W., Li, R., Qie, J., Zhen, B., Wang, Y., et al. (2018). A proteomics landscape of circadian clock in mouse liver. Nat Commun 9, 1553.

Yang, Q., Pando, B.F., Dong, G., Golden, S.S., and van Oudenaarden, A. (2010). Circadian gating of the cell cycle revealed in single cyanobacterial cells. Science 327, 1522–1526.

Yuan, K., and O’Farrell, P.H. (2015). Cyclin B3 is a mitotic cyclin that promotes the metaphase-anaphase transition. Curr. Biol. 25, 811–816.

Zitouni, S., Francia, M.E., Leal, F., Gouveia, S.M., Nabais, C., Duarte, P., Gilberto, S., Brito, D., Moyer, T., Kandels-Lewis, S., et al. (2016). CDK1 Prevents Unscheduled pLk4-STIL Complex Assembly in Centriole Biogenesis. Current Biology 26, 1127–1137.

